# Primary carbohydrate metabolism genes participate in heat stress memory at the shoot apical meristem of *Arabidopsis thaliana*

**DOI:** 10.1101/2020.08.20.258939

**Authors:** Justyna Jadwiga Olas, Federico Apelt, Maria Grazia Annunziata, Sarah Isabel Richard, Saurabh Gupta, Friedrich Kragler, Salma Balazadeh, Bernd Mueller-Roeber

## Abstract

Although we have a good understanding of the development of shoot apical meristems (SAM) in higher plants, and the function of the stem cells (SCs) embedded in the SAM, there is surprisingly little known of its molecular responses to abiotic stresses. Here, we show that the SAM of *Arabidopsis thaliana* senses heat stress (HS) and retains an autonomous molecular memory of a previous non-lethal HS, allowing the SAM to regain growth after exposure to an otherwise lethal HS several days later. Using RNA-seq, we identified genes participating in establishing a SAM-specific HS memory. The genes include *HEAT SHOCK TRANSCRIPTION FACTORs* (*HSFs*), of which *HSFA2* is essential, but not sufficient, for full HS memory in the SAM, the SC regulators *CLAVATA1* (*CLV1*) and *CLV3*, and several primary carbohydrate metabolism genes, including *FRUCTOSE-BISPHOSPHATE ALDOLASE 6* (*FBA6*). We found that expression of *FBA6* during HS at the SAM complements that of *FBA8* in the same organ. Furthermore, we show that sugar availability at the SAM is essential for survival at high-temperature HS. Collectively, plants have evolved a sophisticated protection mechanism to maintain SCs and, hence, their capacity to re-initiate shoot growth after stress release.

## Introduction

The shoot apical meristem (SAM) is a highly organized tissue that is essential for proper above-ground growth of plants (1). Descendants of a small number of stem cells (SCs) in the central zone of the SAM form shoot structures like leaves, flowers and derivatives thereof (seeds and fruits). The SC population has self-maintaining and self-renewal capacities that allow plants to develop new organs throughout their entire lifespan (2). SC homeostasis is maintained by a negative feedback loop involving CLAVATA1 (CLV1), CLAVATA3 (CLV3), and WUSCHEL (WUS). WUS protein promotes SC identity by inducing expression of the *CLV3* gene, while the CLV3 peptide interacts with the receptor-like kinase CLV1, thereby suppressing *WUS* expression and hence SC proliferation (3). Since the growth and initiation of new organs depend on SC activity, perturbation of the WUS-CLV control module affects the plant’s architectural organization (1). As cells of the SAM cannot photosynthesize, due to a lack of functional chloroplasts, they are heterotrophic and therefore depend on sugar supply from photosynthetically active sources such as cotyledons and leaves (4). Given its biological importance for seedling survival and shoot growth, the SAM is presumably fortified in particular ways against diverse environmental stresses the plant may encounter.

Although the core regulatory mechanisms that control SAM formation and SCs’ functions have been extensively addressed over the last two decades (2), we have little insights into responses of the SAM and SCs to environmental stresses and how their formation and function are maintained under adverse environmental conditions. Recent studies have shown that the *Arabidopsis thaliana* (Arabidopsis) SAM can sense and adaptatively respond to changing soil nitrate levels (5) as well as to carbon depletion (6). Both observations support the notion that the SAM has the competence to sense and adapt to environmental stresses to maintain shoot growth.

Besides mechanisms enabling acute responses to stress (7, 8), plants have evolved a stress adaptation system, called stress memory, that ‘primes’ (or prepares) them to survive a severe stress that follows a moderate stress which occurred days before. The stress memory supports plants to store information about the previous stress during a ‘memory’ phase to better prepare for a subsequent, potentially more severe (triggering) stress event (9). This information or ‘memory’ of the exposure to the priming stress allows plants to survive an otherwise lethal stress occurring days later (10, 11). The transcriptional memory of a previous stress includes two categories: (i) genes whose expression is changed by the first stress and show a sustained change of expression during the memory phase, or (ii) genes that show hyper-induction upon a recurrent (triggering) stress allowing them to faster and/or strongly respond to a second stress (12). One of the most well-documented memory phenomena is cold acclimation, which involves coordinated transcriptional, metabolic, and physiological responses to sub-optimal temperatures that increase the plants’ resistance to subsequent ‘colder’ temperatures (13, 14). Such a perception mechanism has clear adaptive value in continually fluctuating natural environments; a cold day is more likely to be followed by another cold day (or even colder day) than a warm day. Molecular mechanisms involved in cold stress memory are well understood, and recently some progress has been made in unraveling transcriptional memory of heat stress (HS) in whole Arabidopsis seedlings (9, 11).

Similar to cold stress, HS is a major abiotic factor that dramatically limits plant growth, development, and, consequently, seed production (15). It induces, among others, the expression of genes encoding (*inter alia*) various types of HEAT SHOCK PROTEINs (HSPs), which confer thermotolerance by acting as chaperones that facilitate proper protein folding and function (16). Production of HSPs is induced by HS in all higher organisms and is an energy-costly process controlled by HEAT STRESS TRANSCRIPTION FACTORs (HSFs) (17). In Arabidopsis, there are 21 known HSFs, grouped into three main classes (A, B, C), based on structural differences (17). Importantly, not all *HSFs* are HS-inducible. Depending on the stress type, each factor controls a specific regulatory network. While class A HSFs are positive regulators of the HS response, members of class B act as repressors (18). HSFs bind to heat shock elements (HSEs, with a conserved 5’-nGAAnnTTCn-3’ sequence) in promoters of HS-inducible genes (17). To date, only three of the 21 HSFs (HSFA2, HSFA1a, and HSFA1e) have been demonstrated to play a role in HS memory in Arabidopsis. The *hsfa2* mutant is defective in thermomemory, and HSFA2 protein is required for maintenance of high expression of several HS-memory-related genes, but not for their initial induction (19, 20).

In summary, the SAM and SCs are essential for plants’ shoot growth (21), and thermoprimed plants can grow after otherwise lethal HS (10, 11), but the mechanisms involved in priming-induced protection of the SAM are unknown. It is not even known whether the SAM can generate an HS memory autonomously, and, if so, whether the molecular mechanisms differ from those of other organs. Also it is conceivable that the SAM generates a non-autonomous HS memory depending on signals and/or metabolites from sensing cotyledons or young leaves that protect the SAM from exposure to stress. Here, we show that the SAM, including its SC population, can generate a strong autonomous HS transcriptional memory with primary carbohydrate metabolism, protein folding, and meristem maintenance genes acting as HS memory components at the SAM. We demonstrate that priming promotes meristem maintenance from otherwise lethal HS. Moreover, we show that sugar availability is a crucial factor for thermomemory. Finally, we demonstrate that HSFA2 is an important, but not sufficient, transcriptional regulator of thermomemory in the SAM, suggesting that a distinct and complex regulatory network governs HS responses in the tissue.

## Results

### Thermoprimed plants fully recover shoot growth after a second heat stress

Since the SAM is responsible for overall shoot growth, we investigated how thermopriming affects plant growth and development under both long-day (16h light/8h dark) and day-neutral (12h light/12h dark) photoperiods. For this, we subjected five-day-old vegetative seedlings of Arabidopsis Col-0 plants to an established thermomemory assay (Fig. 1*A*) (10). Primed and triggered (PT) plants remained green and continued to grow in both photoperiods (Fig. 1 *B* and *C*), as previously reported. In contrast, unprimed seedlings subjected solely to the HS trigger (T seedlings) gradually collapsed and died (Fig. *1B* and *C*). Analysis of growth behavior, using a 3D time-lapse imaging system (Fig. 1*D-F* and *SI Appendix*, Fig. S1*A* and *B*) (22, 23), revealed that the total rosette area of PT plants exponentially increased between days 4 to 16 after priming (DAP; 1-13 days after triggering; DAT). We found the same for control (C) and moderate heat-primed (P) plants not exposed to a second triggering HS (Fig. 1*D*). This observation confirmed the importance of priming for the ability of Arabidopsis plants to survive a subsequent triggering stimulus. Importantly, however, non-treated C plants had significantly larger (four-fold, *P*≤ 0.001) rosette areas than PT plants at the end of the imaging period, indicating that the PT treatment caused an approximately one-week delay in growth (Fig. 1*D*). The relative rosette expansion growth rate (RER) was significantly reduced in PT plants for several days after the triggering but started to recover at around 7 DAP (4 DAT) (Fig. 1 *E* and *F*), indicating that their lower rosette area at the end of the experiment was due to growth inhibition immediately after the triggering and that growth rate fully recovered a few days later.

**Fig. 1.**
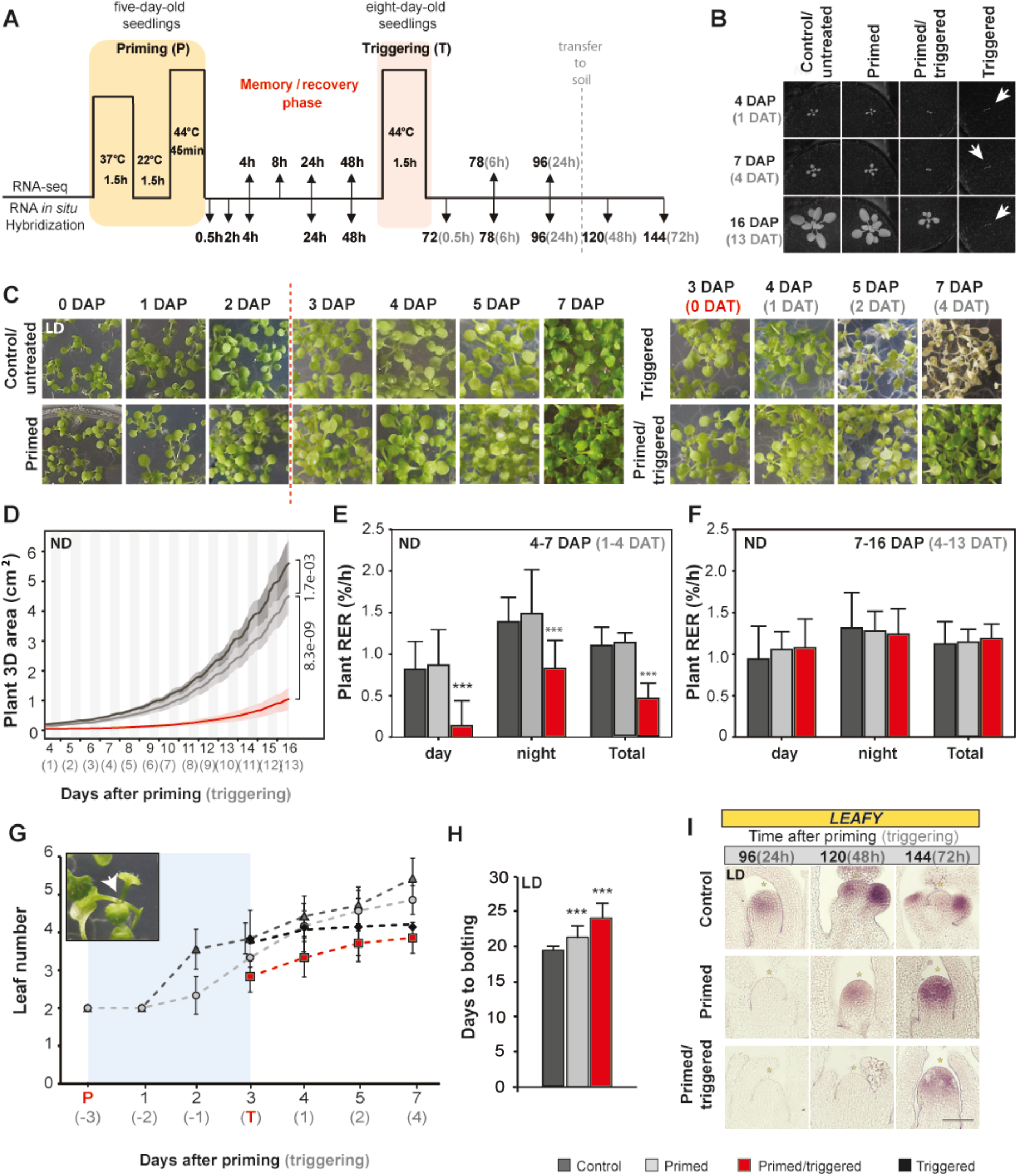
Growth and development of thermoprimed Col-0 seedlings in neutral day (ND) and long day (LD) conditions fully recovers after the treatment. *(A)* Schematic representation of the experimental set-up. Five-day-old seedlings grown in MS media with 1% sucrose were subjected to a moderate priming HS at 6h after dawn, followed by a three-days memory/recovery phase, and then subjected to a second triggering HS at 9h after dawn. One day after triggering (DAT), seedlings were transferred to soil to monitor growth and development. Samples were taken at different time points for RNA-seq analysis or RNA *in situ* hybridization. *(B-C)* Images of Col-0 plants during and after priming (DAP, days after priming) and triggering treatments. Images in panels *(B)* and *(C)* were used for measuring growth behavior after treatment in ND or LD condition, respectively. *(D)* Increase of total plant 3D area over time of control (C), primed (P), and primed and triggered (PT) plants (*n*≥6 for each condition). *(E-F)* Comparison of mean relative expansion growth rate (RER, % per h) of control C, P, and PT plants analyzed after the thermopriming treatment during day, night, and in total for 4-7 DAP or 7-16 DAP. The data are calculated from the plot shown in *(D). (G)* Leaf initiation rate of Col-0 plants grown in LD conditions (*n*>10). The memory/recovery phase is marked in blue. *(H)* Flowering time based on ‘days to bolting’ in LD conditions (*n*=20). *(I)* RNA *in situ* hybridization using a *LEAFY* antisense probe on longitudinal sections through apices of C, P, and PT plants in LD conditions. Time is given in hours (h) after priming (black color) and triggering (grey color) treatments. Error bars indicate s.d.; asterisks indicate a statistically significant difference (Student’s *t*-test: \*\**P*≤ 0.01; \*\*\**P*≤ 0.001) from the control conditions *(D-H)* or meristem summit *(I)*. Scale bar, 100µm *(I)*. See also *SI Appendix*, Fig. S1, and Table S1.

PT plants may develop a smaller rosette area than C plants because of a reduced leaf initiation rate (LIR). We tested this possibility by monitoring leaf emergence (Fig. 1*G* and *SI Appendix*, Fig. S1*C*), which showed that PT treatment reduces LIR. Importantly, however, LIR remained unabated in P plants. After a short delay following the triggering event it resumed in PT plants but entirely ceased at ∼2 DAT in T plants (see insert in Fig. 1*G*).

Since P and PT plants flowered 2 and 5 days, respectively, later than C plants (Fig. 1*H* and *SI Appendix*, Fig. S1*D* and Table S1), we performed toluidine blue staining of longitudinal sections through meristems, and RNA *in situ* hybridization using floral marker transcript *LEAFY* (24). This allowed us to determine the exact time of the floral transition of Col-0 plants. As shown in Fig. 1*I* and *SI Appendix*, Fig. S1*F*, P plants initiated floral transition a day later than C plants. In contrast, flower formation was delayed by approximately 2 days in PT plants, demonstrating that the SAM of C, P and PT plants all remained in the vegetative stage during the thermopriming (i.e., they had not induced flowering). Hence, the transcriptome changes induced by priming (see following chapter) are not caused by a shift in development phase between C and P or PT plants. In summary, P plants survived when the triggering stimulus was applied, with only a temporary reduction of shoot growth. In contrast, in unprimed plants the triggering stress damaged the existing leaves, blocked the formation of new leaves at the SAM, and limited further shoot development. Thus, the priming treatment induces mechanisms that protect the SAM from the growth-terminating damage observed in T plants.

### Identification of HS memory genes in the SAM

Given that transcriptome changes in the SAM are not induced by developmental transitions (see above), and that the SAM and SCs have a crucial role in shoot regrowth after a priming HS, we investigated how the SAM responds to changes in HS treatments by performing RNA-sequencing (RNA-seq). To this end, we manually dissected and analyzed the transcriptome of shoot apices containing young leaf primordia from P plants and non-treated C plants at selected time points after moderate heat priming (4, 8, 24, 48 and 78h into the recovery/memory phase), and at 6 and 24h after a second exposure to the triggering HS from C, P, PT, and T plants (Fig. 1*A* and *SI Appendix*, Fig. 2 and Data S1). Principal Component Analysis (PCA) of the transcriptome data clustered the shoot apex samples into three separate groups (Fig. 2*A* and *SI Appendix*, Fig. 2). As expected, one group included all samples of C apices (C4, C8, C24, C48, C78, and C96, *i*.*e*., apices of untreated C plants collected 4, 8, 24, 48, 78, and 96h after priming). Interestingly, this cluster also included the P apices harvested 24, 48, and 78h after priming (P24, P48, and P78 samples) and PT apices exposed to both stimuli harvested 96h after priming and, thus, 24h after triggering (PT96 samples) were assigned to the same group (Fig. 2*A*). The clustering of P24, P48, P78, and PT96 together with C samples suggests that gene expression patterns in the SAM of P and PT plants are rapidly reset (within 24h) for most genes to control-like patterns after priming and triggering treatments; thus, the resetting is faster in the shoot apex than in whole seedlings following an identical treatment, where resetting reportedly takes longer than 24h (25). The two other groups included the P4 and P8 primed samples, and triggered-like (PT78, T78, and T96) samples. As expected, after the exposure to heat priming, hundreds of genes were significantly differentially expressed (DE) between apices of P and C plants (Fig. 2*B*; Table S2): 1,175 genes at 4h, 780 genes at 8h, and 203 genes at 48h. At 78h, no significant differences in gene expression between shoot apex samples of P and C plants were observed. Thus, the shoot apex of Col-0 plants very rapidly senses temperature changes but also resets relatively fast after priming to control levels.

**Fig. 2.**
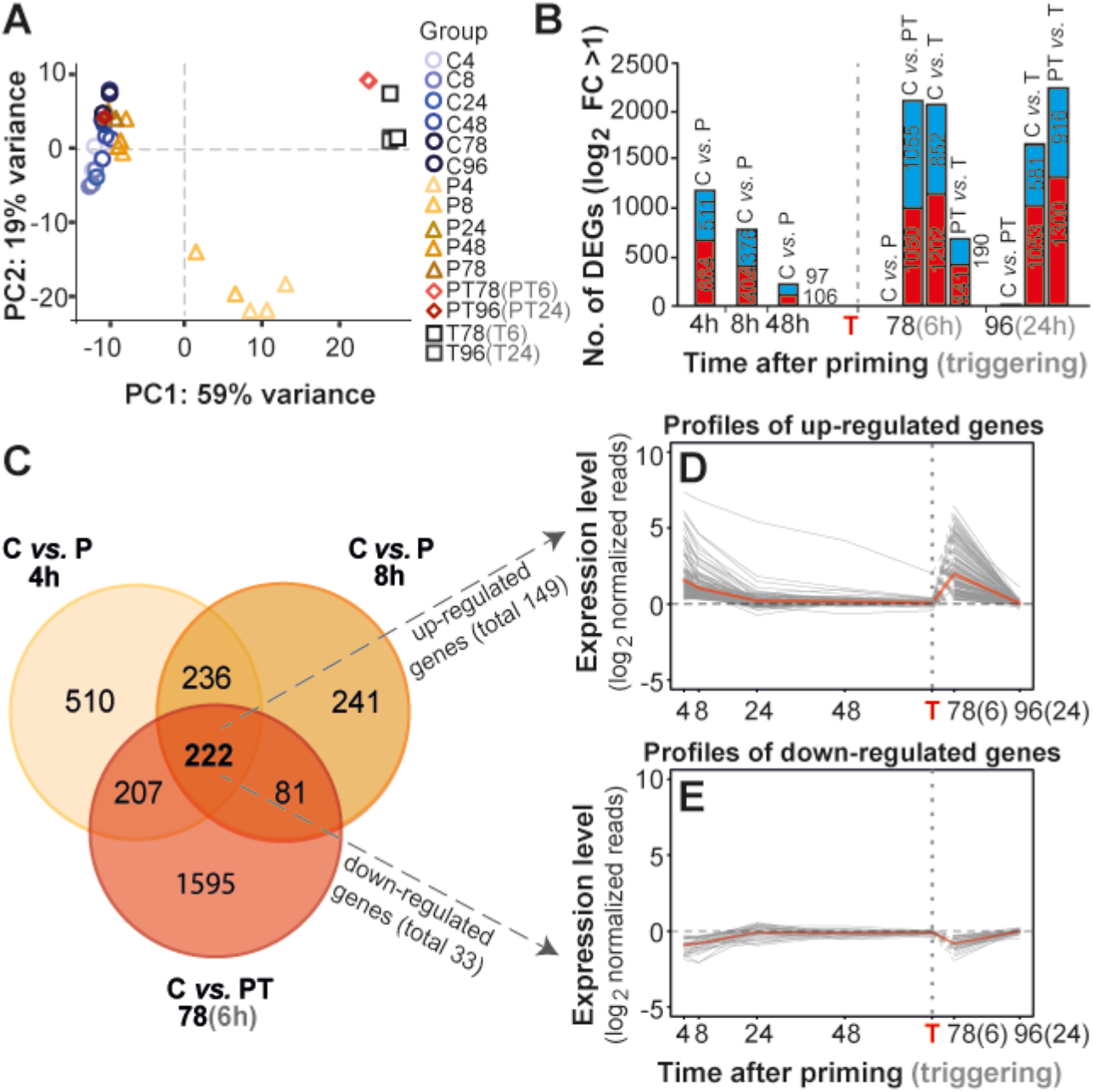
Identification of thermomemory-associated genes in the shoot apex. *(A)* Clustering of the relationship between meristem samples of control (C), primed (P; triangles), primed and triggered (PT; diamonds), and triggered (T; squares) plants during the thermomemory treatment by principal component analysis (PCA) forming three groups. *(B)* Total number of differentially expressed genes (DEGs) between the samples at different time points (in brackets are the numbers of up- (red) or down-regulated (blue) genes; for details, see Methods). *(C)* Venn diagram of DEGs at 4h and 8h after priming and 6h after triggering compared to the control. The overlap represents memory genes at the SAM of Col-0 plants during thermopriming. *(D-E)* Profiles of consistently up-regulated *(D)* and consistently down-regulated *(E)* genes at 4h, 8h, and 78h (6h after triggering) after priming compared to control plants. Profiles were calculated by subtracting the normalized expression values of untreated control plants from the normalized expression values of P or PT plants. The bold red line represents the average profile. The vertical dashed line represents the time point of triggering (T) treatment. Time is given in hours (h) after priming (black color) and triggering (grey color) treatments. Note, that the SAM of only-T plants is hypersensitive to HS, leading to lethality shortly after triggering, therefore, a transcriptomic comparison between PT and T plants (alive *versus* dead tissue) was not performed. See also *SI Appendix*,Fig. S2 and S3, Data S1.

Interestingly, we observed a faster transcriptional response of the shoot apex after triggering in PT plants compared to plants subjected to only the triggering stress. For example, 24h after triggering, expression of only a single gene significantly differed between PT and C samples (Fig. 2*B*; Table S2). Thus, priming enables gene expression to return to control levels within 24 h of triggering, while expression of many genes stays high in unprimed and triggered seedlings. For example, 6-24h after triggering 1,500 – 2,000 genes were significantly DE in T plants, relative to controls, and T plants subsequently died (Fig. 2*B*; Table S2).

We were particularly interested in identifying transcriptional HS memory genes, which, according to the literature (12), are genes showing hyper- or hypo-responsiveness when exposed to the second, more severe stress (memory genes, as defined in *SI Appendix*, Fig. S3*A*). In earlier studies, genes with sustained up- or down-regulation in whole seedlings during the memory phase, without being necessarily DE after a triggering HS, were regarded as thermomemory-associated genes (10, 11). As Arabidopsis does not survive a triggering HS without a priming HS, we were particularly interested in DE genes responding to the triggering treatment after a previous priming treatment. We considered such genes as of particular importance for enhancing HS tolerance and the establishment of HS memory.

As outlined in *SI Appendix* Fig. S3*A*, shoot apex transcriptional HS memory genes were defined as genes that were expressed significantly higher or lower in apices of primed plants 4 and 8h after the priming than in C plants, and higher/lower 6h after the triggering (78h after priming). In total, we identified 394 transcriptional HS memory genes in the shoot apex, of which 217 were upregulated, and 177 were downregulated (Table S2, *SI Appendix*, Fig. S3*B*). Furthermore, to identify high-confidence shoot apex memory genes, we introduced a second criterion, i.e., a fold change in gene expression of |log_2_FC|>1. In total, 149 upregulated and 33 downregulated genes, *i*.*e*., 182 genes, met both criteria (Fig. 2*C-E*; Data S2). Genes induced or downregulated by the priming HS, but not again by the triggering HS after the three-day recovery period, were not regarded as memory genes but recognized as primary stress-responsive genes (11). As expected, several *HSP* family members were among the 149 transcriptional HS memory genes upregulated by the priming and triggering HS treatments in the shoot apex. These include cytosolic *HSP17*.*6A*, nuclear-encoded mitochondrial *HSP22* and chloroplast *HSP21*, as well as five other small *HSPs* (Fig. 3*A*, *SI Appendix*, Fig. S4*A*), suggesting that HS-protective mechanisms are active in all cellular compartments of the SAM. As previous HS studies did not analyze the responses in meristematic tissues, we confirmed the RNA-seq data by RNA *in situ* hybridization using *HSP-* specific (*HSP17*.*6A, HSP21*, and *HSP22*) probes, and by quantitative reverse transcription – polymerase chain reaction (qRT-PCR; Fig. 3*B* and *C* and *SI Appendix*, Fig. S4). We found that expression of the *HSP*s was rapidly induced in all SAM domains, within minutes of a moderate priming HS, then gradually declined until 8h, whereas at 24h after priming *HSP22* was the only detectable *HSP* transcript in the SAM. Moreover, in PT plants we observed hyper-induction of *HSP* genes in the SAM after HS triggering compared to P plants, supporting the RNA-seq results (Fig. 3). These findings demonstrate that *HSP17*.*6A, HSP22*, and *HSP21* are *bona fide* memory genes acting in the SAM. Importantly, we also identified primary carbohydrate metabolism genes involved in sugar metabolism (particularly glycolysis), including *FRUCTOSE BISPHOSPHATE ALDOLASE 6* (*FBA6*), *PYRUVATE KINASE 4* (*PKP4*), and *UDP-GLUCOSE PYROPHOSPHORYLASE 2* (*UGP2*) (Fig. 3*A* and *SI Appendix*, Fig. S4; Data S2), strongly indicating that carbohydrate conversion is essential for the HS memory of the SAM. To obtain information on expression patterns at higher spatial resolution, we selected *FBA6* as a probe for RNA *in situ* hybridization. *FBA6* transcript was barely or not detectable in the SAM of non-stressed plants. However, its transcript abundance increased at 2 and 4h after priming, leading to expression in the organizing center and central, peripheral and rib zones of the SAM (Fig. 3*B*). This result demonstrates differences in the temporal dynamics between the HS memory genes responding at the SAM; for comparison, transcripts of *HSP* memory genes were already induced at 0.5h after the priming. Moreover, expression of *FBA6* at the SAM was even more strongly and faster (already within 0.5h) induced after the triggering HS compared to the priming stimulus. We confirmed the transcriptional induction of *FBA6* and other primary carbohydrate metabolism genes in the SAM of PT plants relative to controls (C plants) by qRT-PCR (Fig. 3*C* and *SI Appendix*, Fig. S4). Thus, *FBA6* is a *bona fide* SAM memory gene in vegetatively growing plants. Importantly, none of the primary carbohydrate metabolism genes were previously reported to be components of the HS memory machinery.

**Fig. 3.**
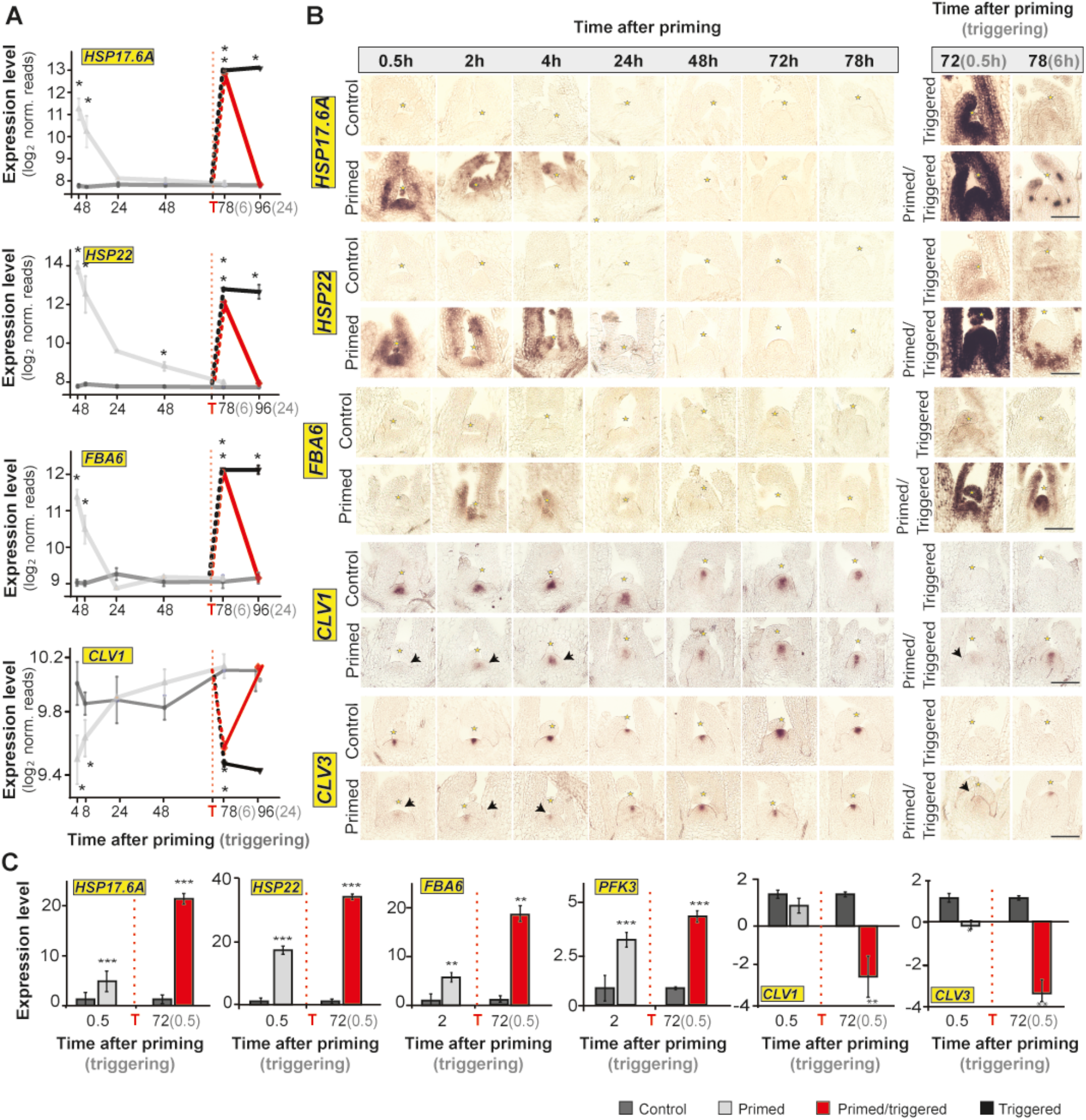
The shoot apical meristem (SAM) displays priming capacity and a transcriptional memory. *(A)* Relative expression level of HS memory genes: *HEAT SHOCK PROTEINs* (*HSP17*.*6A* and *HSP22*), *FRUCTOSE-BISPHOSPHATE ALDOLASE 6* (*FBA6*), and *CLAVATA1* (*CLV1*) at the shoot apex of Col-0 plants during thermopriming, obtained by RNA-seq (*n*=3). *(B)* RNA *in situ* hybridization using *HSP17*.*6A, HSP22, FBA6, CLV1*, and *CLV3* as probes on longitudinal sections through meristems of control, primed, primed and triggered, and triggered plants. Scale bars, 100µm. *(C)* Expression level of *HSP17*.*6A, HSP22, FBA6, PFK3, CLV1*, and *CLV3* at the SAM of Col-0 plants during thermopriming obtained by qRT-PCR (*n*=3). Note that plants were grown in MS media with 1% sucrose. Time is given in hours (h) after priming (black color) and triggering (grey color) treatments. The vertical dashed line represents the time point of triggering (T) treatment. Error bars indicate s.d. (*n*=3). Asterisks indicate a statistically significant difference (RNA-seq, \**P*≤ 0.05 adjusted with Benjamini-Hochberg procedure for multiple testing correction; qRT-PCR, Student’s *t*-test: \*\**P*≤ 0.01; \*\*\**P*≤ 0.001) from the control conditions or meristem summit *(B)*. See also *SI Appendix*, Fig. S4.

Furthermore, among the significantly downregulated memory genes was the SAM-specific leucine-rich repeat receptor-like kinase *CLAVATA1* (*CLV1*) (Fig. 3*A*, *SI Appendix*, Fig. S3*B*; Table S2). RNA *in situ* hybridization and qRT-PCR analyses confirmed that expression of *CLV1* was downregulated in the SAM of P plants relative to controls immediately after priming (Fig. 3*B* and *C*). Notably, *CLV1* transcription was even more downregulated in the SAM of PT plants, revealing a clear memory pattern. Furthermore, expression of *CLV3*, which encodes the CLV1-binding peptide ligand, followed the same type of expression pattern (Fig. 3*B* and *C*). This observation suggests that the SCs directly sense the HS treatments, as genes specifically expressed in the SC responded to the thermopriming.

Taken together, our data suggest that key genes of primary carbohydrate metabolism and responsible for meristem maintenance, as well as genes involved in protein folding and repair, are important for thermomemory in Arabidopsis. Importantly, lack of recovery of *CLV1* and *CLV3* expression in the SAM of triggered, unprimed plants shows that priming protects the SAM from the negative, growth-ceasing impact of an otherwise lethal HS. Hence, we demonstrated that the SAM of T plants is hypersensitive to HS, leading to lethality shortly after triggering.

Next, we searched for commonalities in gene responses between the shoot apex and two previously reported studies of whole seedlings at a time point available for all three datasets (4h after priming, obtained in identical priming setups; *SI Appendix*, Fig. S5) (10, 11). First, we searched for genes up- or downregulated at 4h after priming compared to control by applying the same conservative cutoff to all three datasets (|log_2_FC|> 2). In this way, 2,521 genes were DE in at least one of the three datasets. Importantly, at 4h after priming the shoot apex shared only 119 (of 364) and 295 (of 1,316) DE genes with whole seedlings (10, 11). The intersection of all three datasets at that time point included only a small number of 100 genes. Thus, the majority of the transcripts altered only in the shoot apex (1,045 of 1,359 genes), including *FBA6*, are most likely involved in generating the shoot apex HS memory. We next established a heatmap to present the expression levels of the 2,521 genes regulated at 4h after priming considering time points common to all three datasets, i.e., early (4h) and late (48h/52h) after priming. Importantly, many of the genes up- or downregulated in the shoot apex were not, or – if at all – only marginally, affected in whole seedlings (*SI Appendix*, Fig. 5S*B*). These clear differences in gene expression during thermopriming suggest a tissue-dependent control (the SAM *vs*. leaves) in the regulation of HS memory in plants (*SI Appendix*, Fig. S5; Data S2).

**Fig. 4.**
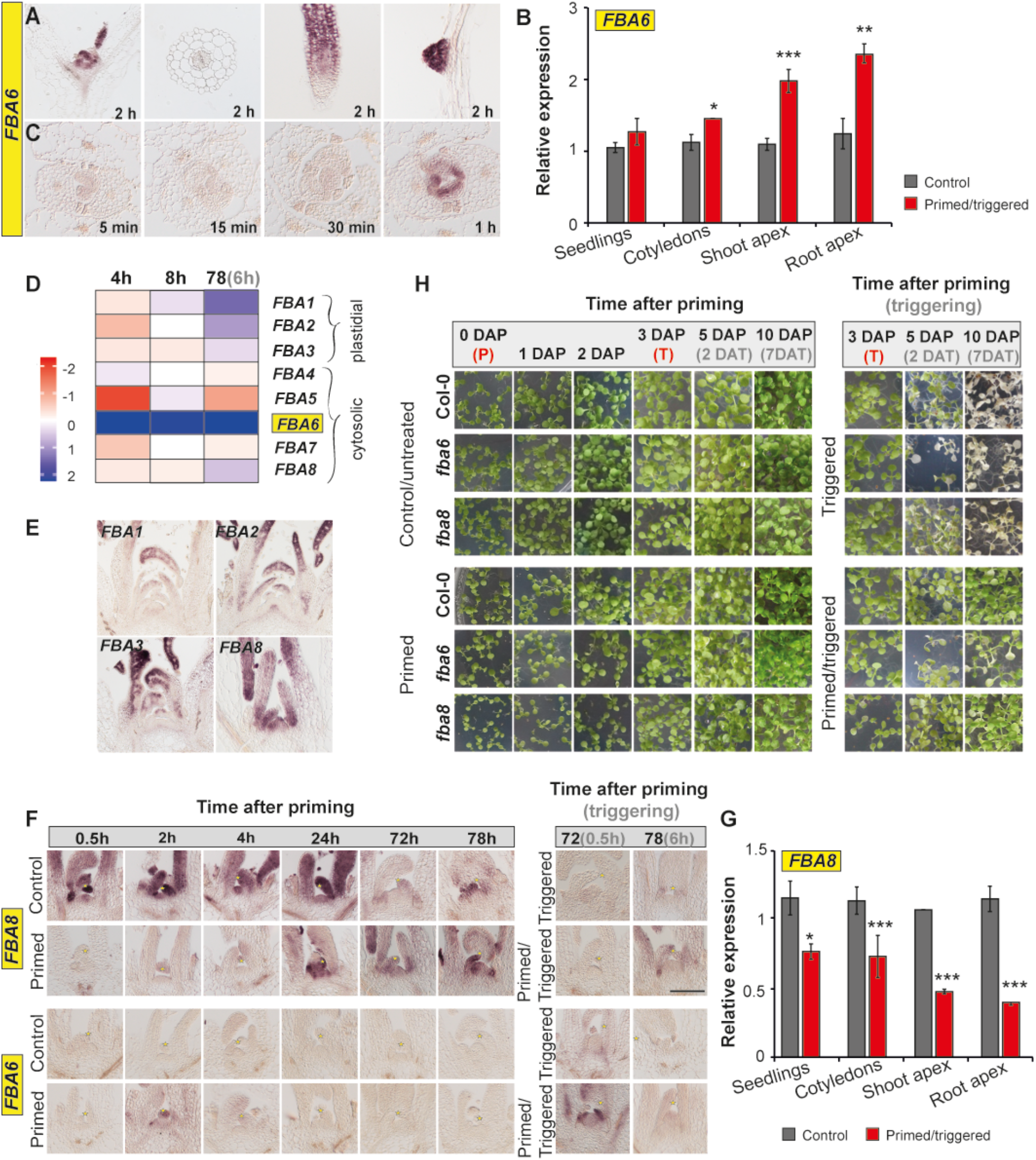
FBA6 affects thermomemory in the shoot apical meristem (SAM). *(A)* Tissue-specific expression of *FBA6* in five-day-old Col-0 plants at 2h after priming. *(B) FBA6* expression in eight-day-old control and primed and triggered Col-0 seedlings at 0.5h after triggering treatment. *(C)* Time-course expression of *FBA6* at the SAM during priming treatment. *(D)* Heat map showing the log_2_ fold change (log_2_ FC) of the expression of all eight *FBA* upregulated (blue) or downregulated (red) genes in Col-0 shoot apex at 4 and 8h after priming and 6h after triggering compared to the control. *(E)* Expression pattern of all identified *FBAs* at the SAM during non-stress conditions. *(F)* RNA *in situ* hybridization using *FBA8* and *FBA6* as probes on longitudinal sections through meristems of Col-0 wild type during thermopriming treatment. Scale bar, 100µm. *(G)* Tissue-specific expression of *FBA8* in eight-day-old control and primed and triggered Col-0 seedlings at 0.5h after triggering treatment. *(H)* Growth recovery phenotype of Col-0, *fba6*, and *fba8* seedlings grown on MS medium with 1% sucrose (Suc) in long-day conditions after priming (DAP) and triggering (DAT) treatments. Error bars indicate s.d. (*n*=3). Asterisks indicate statistically significant difference (Student’s *t*-test: * *P*≤ 0.05; \*\**P*≤ 0.01; \*\*\**P*≤ 0.001) from the control conditions or meristem summit *(G)*. See also *SI Appendix*, Fig. S5.

**Fig. 5.**
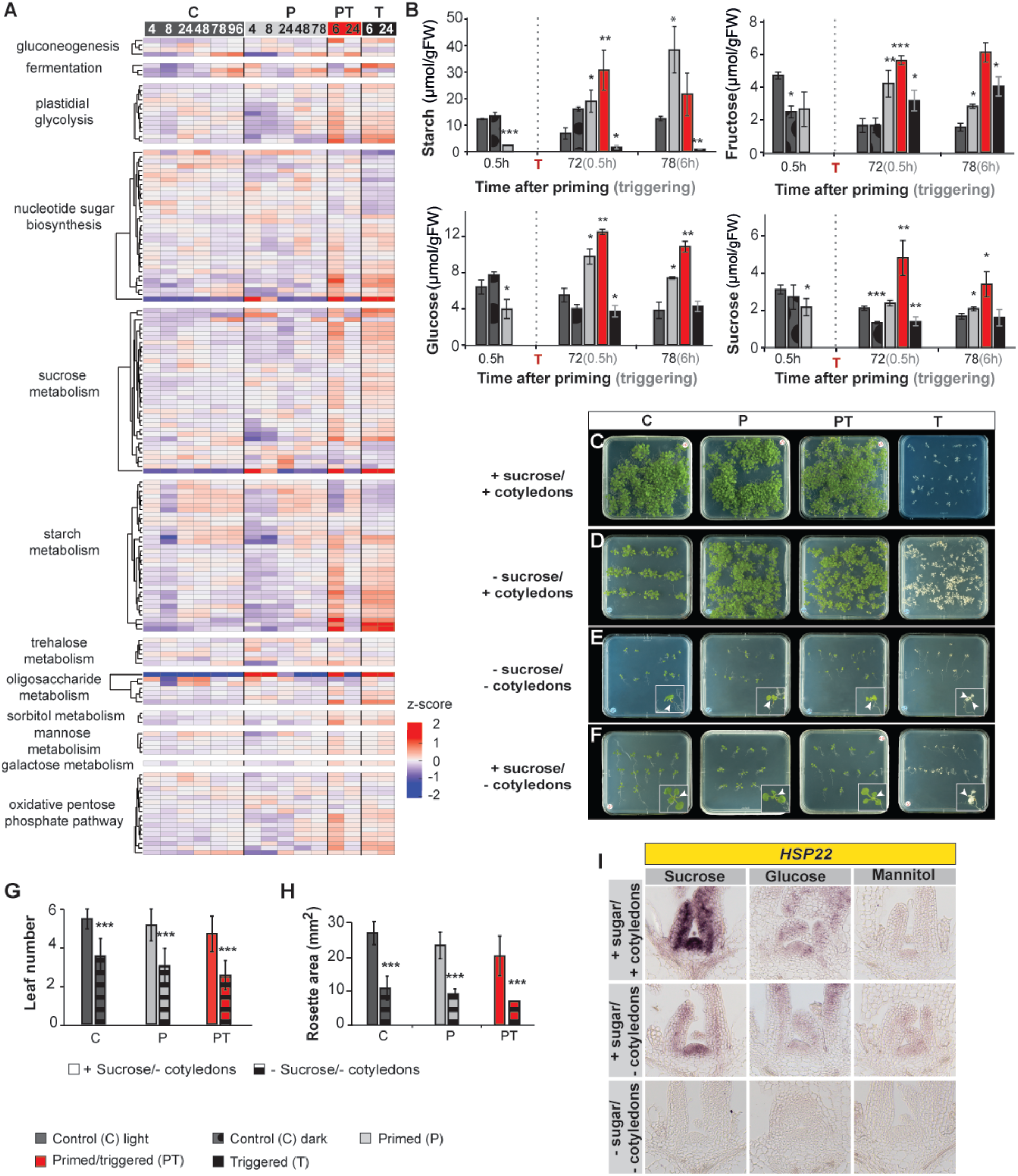
Thermomemory of the SAM is dependent on sugar availability. *(A)* Clustered heat map of differentially expressed upregulated (red) and downregulated (blue) genes (DEGs) of the “Carbohydrate metabolism” category based on level 1 Mapman4 of control (C), primed (P), primed/triggered (PT) and triggered (T) shoot apex samples. Note, that the 6 and 24h time points after triggering in PT and T plants correspond to 78 and 96h after priming in C and P plants. The color scale represents z-score values. *(B)* Soluble sugars and starch content in seedlings during thermopriming. Note that plants were grown in MS media with 1% sucrose. *(C-F)* Thermopriming assay with Col-0 seedlings grown on MS medium with 1% sucrose (Suc; *C*) or without *(D). (E-F)* Col-0 seedlings with detached cotyledons grown on MS medium without *(E)* or with *(F)* 1% sucrose. Note, cotyledons were detached before the priming treatment. Images were taken 10 days after priming (DAP; 7 days after triggering (7 DAT)). *(G-H)* Number of visible leaves *(G)* and rosette area *(H)* in control, primed, and primed and triggered seedlings with detached cotyledons grown with or without 1% sucrose in the medium analyzed at 7 DAT. *(I)* Expression pattern of HS memory gene *HSP22*, determined by RNA *in situ* hybridization on longitudinal sections through meristems of Col-0 wild-type plants grown on sucrose, glucose, mannitol, and no-sugar media, with or without cotyledons. Time is given in hours (h) after priming (black color) and triggering (grey color) treatments. The vertical dashed line represents the time point of triggering (T) treatment. Error bars indicate s.d. (*n*=3). Asterisks indicate statistically significant difference (Student’s *t*-test: \**P*≤ 0.05; \*\**P*≤ 0.01; \*\*\**P*≤ 0.001) from control (light) conditions. See also *SI Appendix*, Fig. S6.

### FBA6 acts as a thermomemory factor in the SAM

Our analysis (Fig. 3; *SI Appendix*, Fig. S5) had revealed that *FBA6* is likely involved in generating HS memory in the shoot apex which contains meristematic tissue. We, therefore, tested whether *FBA6* is a SAM-specific HS memory gene of Col-0 plants during vegetative growth. Tissue-specific localization of *FBA6* expression at 2h after priming revealed that *FBA6* transcript abundance was, in addition to the SAM and young leaf primordia, also elevated in root apical meristems and root primordia, but not in cotyledons, the hypocotyl, or the main root (Fig. 4*A*). This result suggested that FBA6 is specifically generating a HS memory in meristematic, but not other, tissues. Analysis by qRT-PCR confirmed that *FBA6* expression was predominantly induced by heat at the SAM and the root apex (Fig. 4*B*). This demonstrates that HS memory in the non-photosynthetic meristems differs from that in cotyledons. Furthermore, as *FBA6* transcript was detectable at the SAM only after HS we investigated the exact timing of *FBA6* transcriptional activation. RNA *in situ* hybridization revealed that *FBA6* transcript was induced at the SAM approximately 1h after the priming treatment (Fig. 4*C*), and therefore clearly later than the HSP transcripts in the same organ (see Fig. 3).

The Arabidopsis genome encodes eight aldolases (*FBA1-8*). While FBA1 to 3 enzymes are plastid-localized and active in photosynthetic cells, FBA4 to 8 are located in the cytosol and involved in gluconeogenesis and glycolytic carbohydrate degradation (26). Analysis of the shoot apex RNA-seq data (Fig. 4*D*) revealed that only the expression of *FBA6* significantly changed during the priming treatment. Moreover, *FBA6* expression was hyper-induced after HS triggering, suggesting that only FBA6 is involved in generating the SAM’s HS memory. Furthermore, RNA *in situ* hybridization revealed that only *FBA1, 2, 3* and *8* are constitutively operative at the SAM of control/non-treated plants with *FBA8* showing the highest expression (Fig. 4*E*), suggesting that FBA8 is the major aldolase isoform active at the SAM. We, therefore, analysed expression of *FBA8* during thermopriming (Fig. 4*F*) and found a transient downregulation of *FBA8* transcript abundance at the SAM after priming and triggering treatments. Importantly, the downregulation of *FBA8* at the SAM following HS was countered by an induction of *FBA6* (Fig. *4F*), demonstrating that *FBA8* and *FBA6* are oppositely regulated in an HS-dependent manner in this organ. We also found that suppression of *FBA8* by HS was considerably more pronounced in the SAM and the root apex than in whole seedlings and cotyledons (Fig. 4*G*), supporting our finding that HS memory at the SAM is achieved differently than memory in other organs.

We next subjected *fba6* and *fba8* knockout, and Col-0 wild-type seedlings to the thermomemory treatment (Fig. 4*H*). While growth was similar in *fba6* and Col-0 plants during and after priming, growth of *fba8* mutant seedlings was reduced compared to wild-type after priming (*SI Appendix*, Fig. S6*A* and *B*). Both, *fba6* and *fba8* mutants were more sensitive to HS than wild type when subjected to a triggering HS. After priming and triggering, both mutants showed weaker recovery than PT Col-0 seedlings, and a reduced fresh weight (Fig. 4*H*, *SI Appendix*, Fig. S6*A* and *B*), demonstrating that aldolases of primary carbohydrate metabolism are essential components of SAM thermomemory. Lastly, the important role of FBA6 for thermomemory in the SAM was corroborated by the transcriptional response of downstream memory factor *HSP22 (SI Appendix*, Fig. S6*C*). We found that *HSP22* transcript level was strongly compromised at the SAM of the *fba6* mutant compared to Col-0 plants, demonstrating that HS memory at the SAM strongly depends on primary carbohydrate metabolism genes.

In summary, the combined observations provide evidence that primary carbohydrate metabolism plays a crucial role in maintaining HS memory in the Arabidopsis SAM.

### Sugar availability at the SAM is essential for HS memory

The finding that primary carbohydrate metabolism genes including *FBA6* are strongly induced in the photosynthetically inactive SAM, which is photosynthetically inactive, after HS treatment, and the fact that FBAs are key glycolytic enzymes participating in the breakdown of starch or sucrose to generate carbon skeletons and ATP for anabolic processes (27), prompted us to test whether carbohydrate metabolism is altered during HS. First, we generated a heat map to display differences in the expression level of carbohydrate metabolism genes of plants subjected to priming and triggering treatments (Fig. 5*A*). Notable differences were observed between C, P, PT and T plants. In particular, genes involved in plastidial glycolysis, and starch and sucrose metabolism were upregulated after the triggering HS, suggesting that carbohydrate metabolism plays an essential role for survival during temperature stress. To better understand the metabolic differences between the different treatments we subjected C, P, PT and T treated plants to iodine staining to assess their starch content (and hence overall carbohydrate status) after priming and triggering. We found that immediately after priming, starch content declined in primed Col-0 plants compared to control plants (*SI Appendix*, Fig. S7). Interestingly, three days after the priming plants had a higher starch content indicating that more starch accumulated in P plants during the memory phase than in C plants. After triggering, starch content was high in Col-0 PT plants, but low in Col-0 plants subjected to only the triggering HS (T plants).

To further investigate the involvement of sugars in thermomemory, we measured levels of soluble sugars and starch after priming and triggering in Col-0 seedlings (Fig. 5*B*). Confirming the results of the iodine staining and supporting the model that sugars are important for establishing thermomemory, we observed a transient decrease of starch and soluble sugar levels in P seedlings directly after priming, compared to controls, suggesting that carbon consumption is increased in Col-0 plants after the priming HS. Notably, the metabolite levels in P seedlings recovered after three days (72h and 78h after priming) and exceeded significantly those of C seedlings. Furthermore, the levels of those metabolites were even higher in PT than control plants 0.5 and 6h after triggering, demonstrating that carbohydrate metabolism in seedlings responded differently to the second HS than to the first priming HS. Moreover, increased sugar and starch levels were only observed in PT plants, while the levels of these metabolites strongly declined in T plants, similar to P plants subjected to HS priming. These results clearly demonstrate that thermopriming triggers complex changes in starch and sugar metabolism and induces metabolic adjustments in those plants.

To test if establishing thermomemory, and growth recovery after HS, requires energy, we grew Col-0 seedlings on Murashige-Skoog (MS) medium supplemented with (Fig. 5*C*) or without (Fig. 5*D*) 1% sucrose as a carbon source. Additionally, we grew another set of seedlings of which cotyledons were removed before the priming treatment (Fig. 5 *E* and *F*). With or without sucrose in the medium, PT seedlings with cotyledons survived thermopriming and grew further when cultured (Fig. 5*C* and *D*). Seedlings with removed cotyledons also initiated new leaves after priming, irrespective of the presence or absence of sucrose in the medium (Fig. 5 *G* and *H*). However, sucrose was required for seedlings lacking cotyledons to initiate the formation of new leaves and increase leaf size after the triggering HS (Fig. 5 *G* and *H*). This observation demonstrates that plants require carbon to establish thermomemory and restart growth after the triggering HS.

Next, to investigate if the generation of HS memory at the SAM depends on carbon, we analysed the SAM-specific expression of HS memory gene *HSP22* at 0.5h after priming in P plants grown with or without cotyledons on media supplemented with sucrose, glucose, or mannitol. As shown in Figure 5*I*, expression of *HSP22* was enhanced in plants grown with cotyledons and sucrose in the medium. Notably, *HSP22* expression was much lower in plants grown on glucose, and undetectable in plants grown on mannitol, suggesting that HS memory at the SAM is mainly triggered by sucrose. This conclusion is supported by the observation that supply of external sucrose rescues the HS memory in plants lacking cotyledons (Fig. 5*I*). Thus, our results clearly demonstrate that establishing thermomemory in the SAM is a sugar-dependent process.

### HSFA2 is required, but not sufficient, for full HS transcriptional memory in the SAM

Transcription factor HSFA2 is reportedly not involved in the transcriptional activation of HS memory genes in seedlings; rather, it functions in maintaining their enhanced expression after a priming HS (20). To determine whether HSFA2 or other HSFs contribute to the establishment of thermomemory in the SAM, we analyzed the 1-kb 5’-upstream regulatory regions of heat memory genes for basic, and perfect, *cis*-regulatory HSE elements; those might be binding targets of HSFs (28) (Table S3).

We detected basic and perfect HSEs, respectively, in the promoters of 78 and ten high-confidence HS memory genes of the SAM (42.8% and 5.5% of the 182 genes, respectively). This is a highly significant enrichment compared to all Arabidopsis genes (*P* ≤ 0.001, hypergeometric test; for details see Methods), suggesting that these genes are direct targets of SAM HSFs (Table S3 and S4).

Currently, knowledge of the mechanisms underlying binding preferences of HSFs for specific HSEs is missing; we also do not know how many, and which, of the 21 HSFs in Arabidopsis control thermomemory in different tissues (29). However, our RNA-seq data revealed that eight *HSF*s (*HSFA1e, HSFA2, HSFA3, HSFA7a, HSFA7b, HSFB1, HSFB2a*, and *HSFB2b*) might be involved in transcriptional memory in response to thermopriming (Fig. 6*A* and *B*, *SI Appendix*, Fig. S8*A* and *B*), which was confirmed by RNA *in situ* hybridization (*SI Appendix*, Fig. S8*C*). Next, we investigated whether expression of memory genes in the SAM requires HSFA2. First, we confirmed transcriptional memory of *HSFA2* expression in the SAM of Col-0 seedlings by qRT-PCR and RNA *in situ* hybridization (Fig. 6*C* and *D*). We then performed RNA *in situ* hybridization on *hsfa2* knockout mutants, using probes for various memory genes, including genes involved in protein folding (*HSP17*.*6A, HSP21*, and *HSP22*) and primary carbohydrate metabolism (*FBA6*). Expression of these *HSPs* was much weaker in *hsfa2* than in Col-0 wild-type SAMs and, more importantly, there was no detectable induction of the carbohydrate metabolism genes (Fig. 6*E* and *F*; *SI Appendix* Fig. S9).

**Fig. 6.**
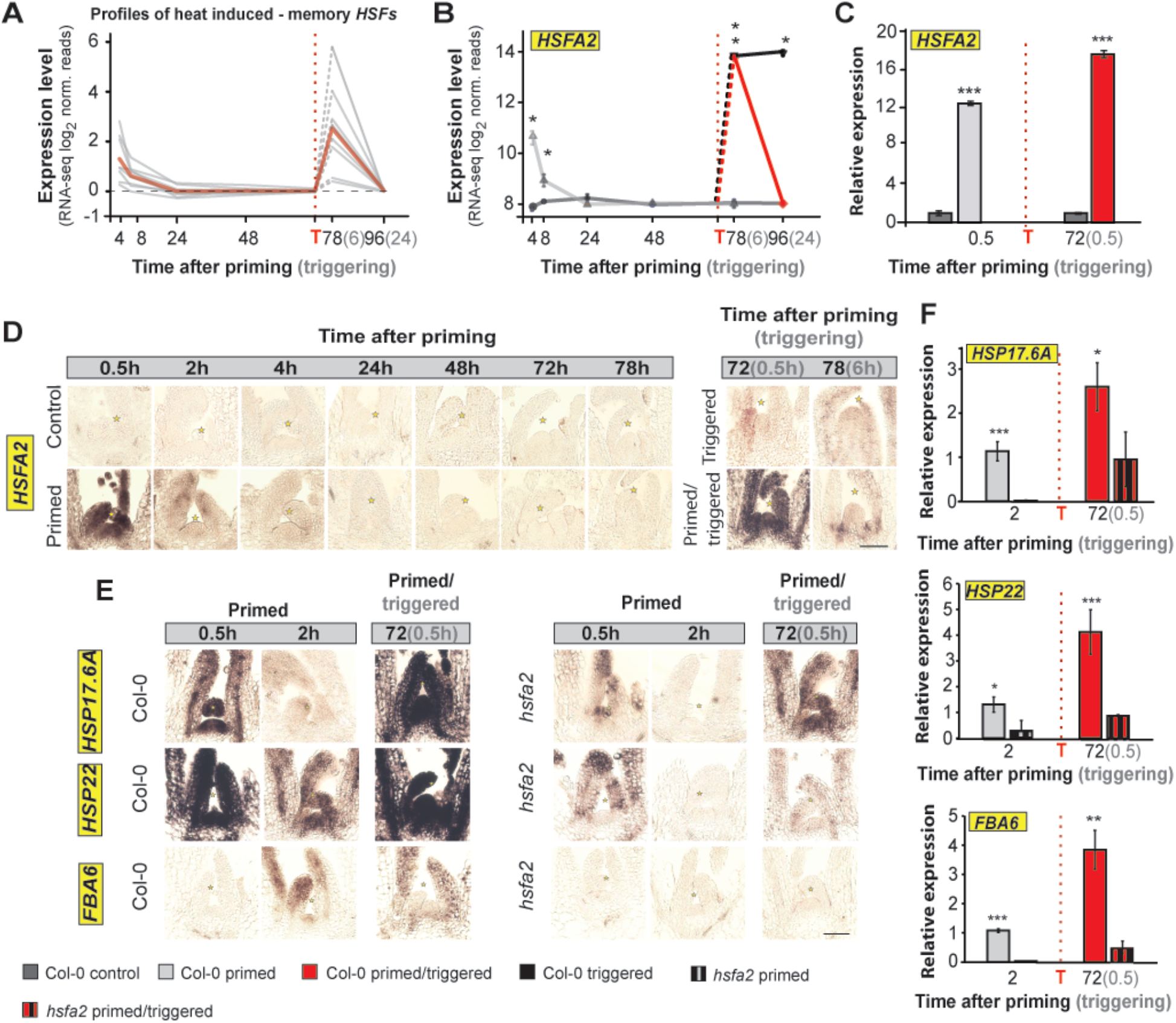
HSFA2 is required, but not sufficient, for full transcriptional memory in the shoot apex. *(A)* Profiles of consistently heat up-regulated *HEAT SHOCK TRANSCRIPTION FACTORs* (*HSF*s) at the SAM of Col-0 seedlings at 4h, 8h, and 78h (6h after triggering) after priming compared to control plants. The bold red line represents the average profile. *(B-C)* Relative expression level of *HSFA2* at the SAM of Col-0 plants during thermopriming obtained by *(B)* RNA-seq and *(C)* qRT-PCR of control plants (dark grey), primed plants (light grey), primed and triggered plants (red), and triggered plants (black) (*n*=3). *(D-E)* RNA *in situ* hybridization using transcript-specific probes for *(D) HSFA2, (E) HEAT SHOCK PROTEIN*s (*HSP17*.*6A* and *HSP22*), and *FRUCTOSE BISPHOSPATE ALDOLASE 6 (FBA6)* on longitudinal sections through meristems of Col-0 and *hsfa2* mutant plants. Scale bars, 100µm. *(F)* Relative expression level of *HSP17*.*6A, HSP22*, and *FBA6* measured at the SAM of Col-0 and *hsfa2* mutant plants during thermopriming. Note that plants were grown in MS media with 1% sucrose. Time is given in hours (h) after priming (black color) and triggering (grey color) treatments. The vertical dashed line represents the time point of triggering (T) treatment. Error bars indicate s.d. (*n*=3). Asterisks indicate meristem summit of statistically significant difference (Student’s *t*-test: \**P*≤ 0.05; \*\**P*≤ 0.01; \*\*\**P*≤ 0.001 *(C, F)* or \**P*≤ 0.05 adjusted with Benjamini-Hochberg procedure for multiple testing correction *(B))* from Col-0. See also *SI Appendix*, Figs. S7 and S9.

These findings demonstrate that in the SAM HSFA2 is required for an initial transcriptional activation of memory genes. This is in stark contrast to previous reports on whole seedlings, which showed that HSFA2 is not involved in the initial transcriptional activation of target genes after priming and required only for maintaining their elevated expression (19, 20). Furthermore, expression of several *HSPs* in the SAM 0.5 h after the priming was weaker than in controls, but not absent in *hsfa2* mutant seedlings, suggesting that HSFA2 is required but is not sufficient for full transcriptional memory in this organ (Fig. 6*E*). This observation is consistent with our finding that seven other *HSFs* apart from *HSFA2* are induced in the SAM during the thermomemory phase.

## Discussion

In Arabidopsis seedlings, a severe (triggering) HS is lethal, while a moderate HS protects from an otherwise lethal triggering HS applied several days later (10, 11). Despite the high academic and commercial interest in understanding how plants survive exposure to high temperature, the molecular mechanisms underlying this phenomenon are not well understood. Although the new above-ground organs formed by plants are initiated by the SAM (30), stress responses of the SAM have rarely been investigated in the past, including those involving thermomemory. To our knowledge, information on responses of the SAM to HS, or its HS memory, is lacking. Most previous studies on HS memory have focused on whole seedlings and their responses to the first moderate priming treatment and the directly following memory phase, neglecting molecular and biochemical responses induced by the second lethal triggering treatment. This observation strongly limits our overall understanding of plants’ responses to recurrent thermostress in their natural habitats.

Our results provide two lines of compelling evidence that the SAM of Arabidopsis directly responds to high-temperature stress. First, leaf initiation (the main developmental read-out of the vegetative SAM) was completely inhibited in T plants, but continued unabated in P plants, and was only delayed in PT plants, demonstrating the importance of priming for the protection of the SAM. Thermoprimed plants grew further after the severe HS triggering, generating new leaves and initiating floral transition, although growth and development were delayed. The inhibitory effect of HS (above 30°C) on growth has been previously reported (31), but the effects of a moderate priming HS on development have not been systematically addressed. We demonstrate here that even a moderate priming HS decreases growth and interferes with development. Moderate priming and severe triggering HS affect growth and development in a manner different from the effect of elevated ambient temperature, which promotes growth and induces earlier flowering in Arabidopsis (32). This observation suggests that delays in growth and transition to flowering in PT plants are adaptive responses that reduce risks of flowering and seed formation during an excessively warm period, and thus potential losses of yield.

Second, we established that the SAM has a distinct thermopriming capacity and transcriptional thermomemory and show that this is a carbon-dependent process. We identified many thermomemory genes whose expression increased or decreased in the SAM after exposure to a moderate, priming HS, with a further up- or downregulation upon exposure to a second severe triggering stimulus. The SAM memory genes included several cytosolic, mitochondrial, and plastidial *HSPs* suggesting that HS-protective mechanisms are active in all cellular compartments of the SAM (10, 11). An unexpected finding of our study, supported by experimental evidence, is that primary carbohydrate metabolism plays a key role in establishing thermomemory in the SAM. First, our analyses identified several primary carbohydrate metabolism genes, including *FBA6*, as *bona fide* thermomemory genes. Among all eight aldolases in Arabidopsis, only *FBA6* showed a hyper-induction in response to HS treatment at the SAM; the transcriptional response of *FBA6* to thermopriming is specific to meristematic tissues including the SAM. Redundancy of genes encoding primary carbohydrate metabolism enzymes is well documented in plants. The different isoforms may provide metabolic robustness, or flexible responses to environmental cues, as seen here for HS memory. Reduced expression of *FBA8* during HS at the SAM was balanced by upregulation of *FBA6* expression in the same organ, indicating that both *FBAs* replace each other’s function during the fluctuating conditions, a behaviour similar to that observed for *NITRATE REDUCTASE 1* and *2* which complement their expression in the SAM (33). Moreover, cytoplasmic aldolases like FBA6 and FBA8 are essential for gluconeogenesis and glycolysis to generate ATP and building blocks for anabolism (27), thereby allowing the cytosolic glycolytic network to provide metabolic flexibility that facilitates acclimation of plants to environmental stresses. Changes in the expression of aldolases seem to be essential for plants during thermopriming, as seen previously for animals, where FBA activity was required for the growth and survival of chronically infected mice (34).

Furthermore, higher plants use sucrose and starch as the principal substrates for glycolysis (27), and our metabolite analysis showed that thermopriming strongly affects carbon reserves in thermoprimed plants. Interestingly, we also noted increased expression of *SUCROSE SYNTHASE 3* (*SUS3*) and *UGP2*, suggesting that sucrose breakdown occurs at that time in the SAM. *SUS*-encoded enzymes catalyse sucrose degradation and play an important role in carbon use in non-photosynthetic cells (35, 36). Moreover, primary carbohydrate metabolism genes such as those identified here as HS memory genes in the SAM are well-established entry points for many key metabolic processes such as glycolysis, a central metabolic pathway for energy production. Our data thus suggest that carbohydrates play a crucial role in thermopriming in the SAM. Sugars are also essential regulators of many developmental and biological processes and important carbon sources for energy metabolism, particularly in heterotrophic organs like the SAM in which no functional chloroplasts exist and which are, therefore, incapable of photosynthesis (4). Intriguingly, sugar (glucose) metabolism and the associated provision of energy also play important roles during stress responses in human and animal SCs (37, 38), suggesting a conserved mode of action of SCs in both photosynthetic and non-photosynthetic organisms. We showed that SCs can directly respond to high temperature, as the meristem maintenance genes *CLV1* and *CLV3* act as HS memory components. Moreover, we demonstrated that priming maintains SC activity and protects the SAM from growth-terminating damage, which otherwise occurs in plants exposed to acute stress.

Next, we demonstrated a clear requirement of sucrose for the establishment of thermomemory in the SAM. The removal of cotyledons before priming impaired thermotolerance of the SAM, which could be restored by an exogenous supply of sucrose. In the presence of sucrose, PT Col-0 seedlings with removed cotyledons grew normal after thermopriming, while the formation of new leaves was significantly reduced in seedlings grown without sucrose and cotyledons. Our observations strongly indicate that cotyledons play an important role for thermomemory by providing sugars to the SAM, harboring the SC population, of stressed plants. Cotyledons are the main photosynthetic sources of fixed carbon in seedlings that provide the energy needed for growth until the first true leaves emerge (39). Lack of cotyledons and sucrose in the medium during thermopriming leads to growth inhibition due to carbon limitation. This observation is in accordance with the weaker expression of the *HSP22* memory gene at the SAM of plants grown without cotyledons but with sucrose, compared to plants grown with both cotyledons and sucrose. The expression differences likely reflect the importance of a clearly defined source-sink relationship between cotyledons (or leaves) and the SAM. In this scenario, sucrose would be more efficiently transported from cotyledons to the SAM than from roots exposed to sucrose in the medium. Cotyledons not only supply phloem-mobile metabolites, but also systemic signals required for full functionality of the SAM. Together, our results strongly suggest that transcriptional induction of *HSP* genes in the SAM requires cellular energy provided by carbon metabolic activity.

An important and most relevant finding of our work is that the SAM, and the SC population it harbors, recruits a HS response (and memory) network that in many aspects differs from that of whole seedlings mainly harboring differentiated cells. Our results clearly demonstrate that a tissue- or organ-specific heat-memory exists in Arabidopsis. Firstly, and importantly, the timing of the expression of several *HSP* memory genes differed between the SAM and whole seedlings. While memory genes in the apex are induced within minutes and remain induced for up to 8h, induction of such genes in whole seedlings occurs after 4-24h, and remains elevated for up to 52h (20), clearly indicating that HS responses are more rapid in the SAM than in most other tissues. We found that *HSFA2*, which was previously reported to be only required for the *maintenance* of thermomemory in whole seedlings (19, 20), is necessary for the *activation* of memory genes at the SAM. *HSFA2* transcript level peaks within 30 min of priming and rapidly declines thereafter, whereas its expression in whole seedlings is reportedly strongest 4h after priming (20). Furthermore, the SAM of the *hsfa2* mutant has largely lost transcriptional memory. As a consequence, the expression of *HSP* memory genes is strongly downregulated shortly after priming in the *hsfa2* SAM. Thus, within the SAM, *HSFA2* is not only responsible for the maintenance of *HSP* expression, which is in contrast to whole seedlings, but also for their initial transcriptional induction. The probable physiological importance of such organ-specific differences in timing and regulation warrants attention in future research. In addition, we showed that different sets of genes, including *FBA6*, are involved in generating HS memory in the SAM and cotyledons. To date, only three of the 21 HSFs in Arabidopsis had been shown to participate in HS memory: HSFA1a, HSFA2, and HSFA1e (19, 20, 29, 40). Here, we provide molecular evidence that eight HSFs are likely involved in thermomemory in the SAM.

Interestingly, although multiple HSFA1 isoforms (a, b, d, and e) are reportedly master regulators of the HS response in Arabidopsis and required for expression of *HSFA2* (41), we only found *HSFA1e* to be transcriptionally induced in the SAM, suggesting that induction of *HSFA2* in the SAM might be only HSFA1e-dependent. Moreover, the glucose-dependent regulation of thermomemory in whole seedlings acts through the HSFA1a isoform (40) whose expression was not affected by priming, or by priming and triggering, stresses in the SAM. This observation provides additional evidence that the mechanisms of thermomemory regulation in the SAM and other organs of seedlings differ.

Our data highlight the complexity of the HS transcriptional memory in the SAM, which depends on carbon metabolism involving (*inter alia*) *FBA6*, and HSF pathways involving HSFA2 as an important, but not the only, transcriptional regulator (Fig. 7). The ability of the SAM, which harbors the key SC population, to respond to environmental stresses and retain ‘memory’ of previous non-lethal stress has clear eco-physiological value. The unique renewal capacity of SCs provides plants with the developmental flexibility required to adjust their developmental processes in response to environmental cues, and to replace body parts lost through damage caused by stresses. This plasticity is particularly crucial for young plants that have not yet initiated axillary meristems or floral transition as their survival depends on a functional SAM and the embedded SCs forming new leaves and flowers.

**Fig. 7.**
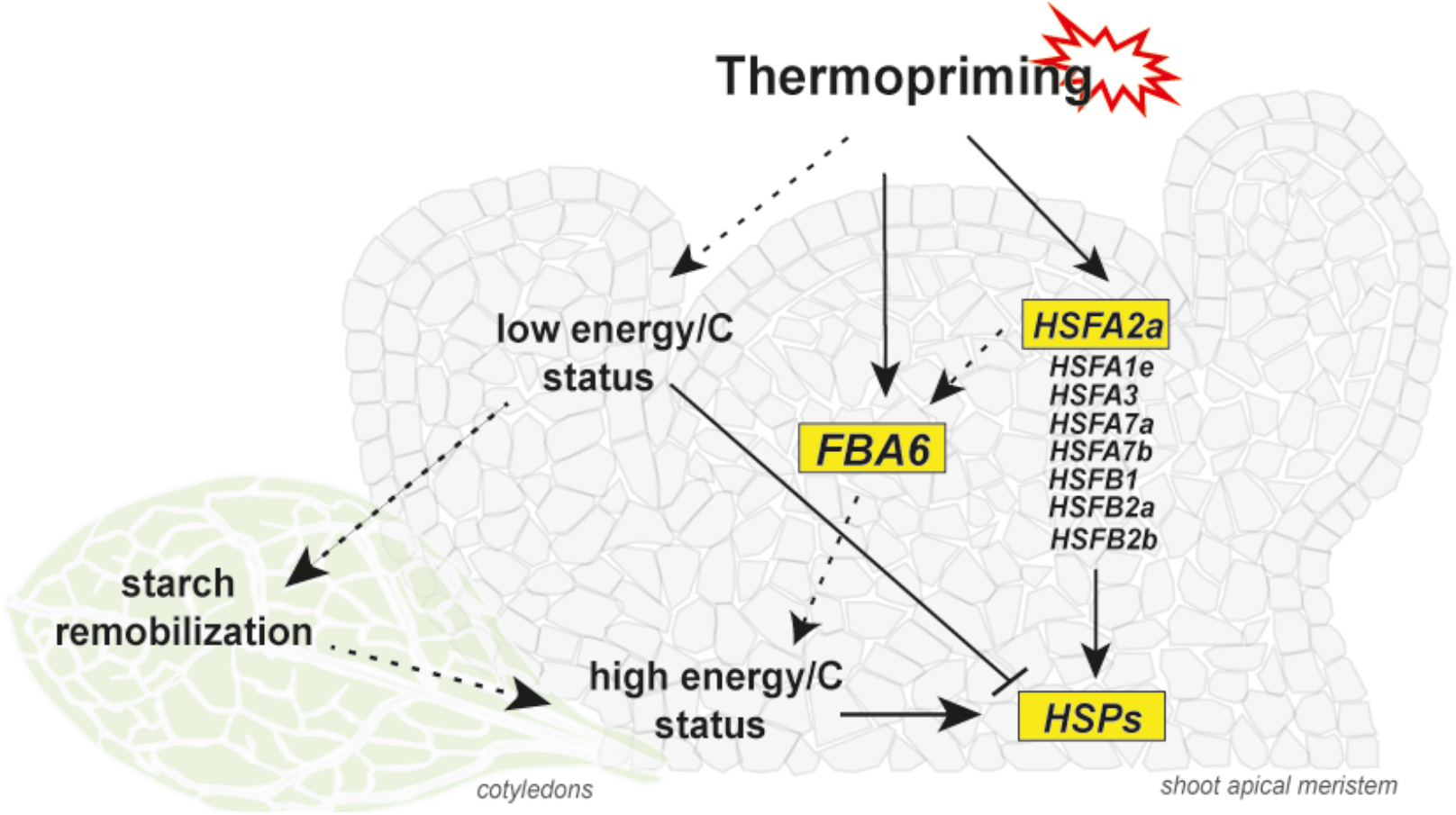
A minimal model for the regulation of heat stress (HS) memory at the shoot apical meristem (SAM). The SAM shows thermopriming capacity and HS transcriptional memory. Thermopriming induces the expression of specific HSFs at the SAM, including the master regulator HSFA2. HSFs might directly bind to HSE in the 5’ upstream regulatory regions of memory genes identified at the SAM. Further, the priming HS affects the sugar availability in plants and activates the expression of primary carbohydrate metabolism genes. Solid lines, direct interactions; dashed lines, indirect interactions.

## Material and methods

Detail description of the material and methods used in this study is described in the supplementary *Appendix*.

### Data availability

The sequencing data sets are available at the NCBI Sequencing Read Archive (SRA), BioProject ID PRJNA505602.

## Acknowledgments

Bernd Mueller-Roeber thanks the Deutsche Forschungsgemeinschaft, Germany (DFG), for funding project A5 within the Collaborative Research Centre 973 ‘Priming and Memory of Organismic Responses to Stress’ (www.sfb973.de). Fritz Kragler thanks the European Research Council (ERC) for funding project Syg Project 810131 (PLAMORF). We thank Eike Kamann from the University of Potsdam (Germany) for cloning work and technical assistance, and Svenja Reeck from the same university for general lab work. We thank Prof. Dr. Dr. h.c. Mark Stitt from the Max Planck Institute of Molecular Plant Physiology, Potsdam, Germany, for constructive comments on the research, and Dr. Lei Yang from the same institute for assisting with taking plate images.

## Author contributions

The research and funding is based on founding observations made by S.B. and B.M.-R. J.J.O. and B.M.-R. conceived the details of the study and designed the experiments involving suggestions made by F.K. J.J.O. carried out the experiments and analyzed the data. F.A. performed growth measurements and analyzed the RNA-seq data, except for generating the cluster heat map (S.G.). M.G.A. measured and analyzed the metabolite data. S.I.R. assisted in performing experiments. J.J.O. and B.M.-R. wrote the manuscript, which was improved considering comments by all authors, who then accepted the final manuscript.

## Supplementary Information Appendix

### Materials and Methods

#### Plant material and growth conditions

*Arabidopsis thaliana* seedlings (ecotype Col-0) were grown in 0.5 Murashige and Skoog (MS) agar media with or without 1% sucrose (w/v) under long-day (LD; 16h light/8h darkness) or neutral-day (ND; 12h light/12h darkness) conditions at 22°C with a photosynthetically active radiation of 160 µmol m^−2^s^−1^. The thermomemory protocol was performed as reported (1). Briefly, 5-day-old seedlings were subjected to priming stimulus at 6h after dawn (1.5h at 37°C; recovery at 22°C for 1.5h; 45min at 44°C), afterwards returned to normal growth conditions (22°C) for 3 days, and then subjected to the triggering treatment (1.5h at 44°C). All thermopriming treatments were performed in a water bath. Seedlings were grown in agar plates until one day after triggering (DAT; 4DAP, days after priming); afterwards, plants were transferred to soil to monitor the growth and development. The *hsfa2-1* mutants were previously reported (2). The *fba6* (SAIL_882_C03) and *fba8* mutants were obtained from the NASC collection, and homozygous lines were confirmed by PCR using the primers presented in Table S5.

#### Growth analysis

Plant rosette area and relative expansion growth rate (RER) of control (unprimed; C; *n*=8), primed (P; *n*=10), primed and triggered (PT; *n*=6), and triggered (T; *n*=10) Col-0 plants grown in ND conditions were analyzed using an established three-dimensional camera-based imaging system with high accuracy and time resolution (3, 4). Briefly, plants were continuously imaged using noninvasive near-infrared light in a growth chamber (model E-36L; Percival Scientific; http://www.percival-scientific.com/), starting one day after triggering (DAT) with photosynthetically active radiation of 160 µmol m^−2^ s^−1^ at the plant level.

In LD conditions, the rosette area of C, P, PT, and T plants (*n*≥15) was determined using the Fiji platform for biological-image analysis (5). The leaf initiation rate (LIR) was analyzed by counting the number of leaves produced by plants every day at the same time point. Additionally, for plants grown at LD, the LIR was determined by dividing the total leaf number (TLN) by the days to bolting (DTB).

#### Flowering time analysis

Flowering time was defined by (i) ‘days to bolting’ (DTB), which is the day on which the first flower bud was visible after germination and the main stem had bolted to 0.5 cm, and (ii) by ‘total leaf number’ (TLN) (see Table S1).

#### RNA extraction and RNA-sequencing (RNA-seq)

Total RNA was isolated from three biological replicates, each containing more than 60 hand-dissected SAMs, using the Qiagen RNeasy Mini kit (Qiagen, Hilden, Germany) or the mirVana™ miRNA Isolation Kit (Invitrogen/Life Technologies, Darmstadt, Germany). Shoot apices were collected 4h, 8h, 24h, 48h, and 78h after the priming (P plants), from control plants (unprimed; C) at the same time points, and from C, P, triggered (unprimed; T), and primed and triggered (PT) plants at 6h and 24h after the triggering. The time points 6h and 24h after triggering correspond to 78h and 96h after priming, respectively.

Library preparation and sequencing were performed by LGC Genomics (Berlin, Germany); Illumina NextSeq 500 V2 was used to generate 75-bp single reads with an average number of ≥100 million reads per sample (Data S1). The adapter-clipped reads were filtered for rRNA and organelle sequences using SortMeRNA (version 2.1b) (6). We used STAR (version 2.5.2b) to align the reads to the TAIR10 annotation of the genome of *Arabidopsis thaliana* and counted the reads per gene using HTSeq (version 0.9.1) (7). Generally, more than 80% of the reads could be uniquely matched to the annotated genes (Data S1). Subsequent analysis of the count data was performed in R (version 3.5.1) (8). The data were normalized by applying variance stabilizing transformation (VST) using DESeq2 (version 1.20.0) (9) for expression pattern plotting. Euclidean distance and Pearson correlation were pairwise calculated between the normalized samples identifying four outlier samples that were filtered (for details see *SI Appendix*, Fig. S2). Furthermore, to increase the power of the subsequent differential gene expression (DE) analysis, for each triplicate we filtered samples whose average Euclidean distance to the remaining two triplicates was more than 50% higher as the distance of the other two replicates to each other, resulting in three additional filtered samples. Thus, the filtered dataset contains 38 samples from 15 different experimental conditions having triplicates or duplicates, except for 24h after priming (P24), which remains a single sample due to the filtering and makes DE analysis including time point P24 not feasible; however, it allows representation without standard deviation in time-course plots (see, e.g., Figs. 3 and 4 and *SI Appendix*, Fig. S4, S6, and S7). DE analysis was performed using DESeq2 and edgeR (version 3.22.3) (10, 11) with the criteria of a ≥ 2-fold up-/down-regulation with an adjusted *P*-value (using Benjamini-Hochberg procedure for multiple testing correction) of less than 0.05 for both methods. The clustered heat maps were generated using DESeq-normalized expression counts of differentially expressed genes (DEGs) belonging to ‘Carbohydrate metabolism’, based on level 1 Mapman4 annotations and were plotted using the ComplexHeatmap package (12). The level 1 annotations were further classified into respective level 2 annotations.

#### Identification of hyper-induced memory genes

All 182 high-confidence memory genes were analyzed for hyper-induction by testing if the expression level of gene X in PT plants was significantly higher (for up-regulated memory genes) or significantly lower (for down-regulated memory genes) after triggering (78h) compared to the expression level of the same genes in primed plants after priming (P4). The expression levels for the treated plants were normalized by subtracting the mean expression value from control plants for the corresponding time points. Statistical tests were performed using two-tailed, two-sample equal variance Student’s *t*-test considering *P*-values ≤0.05 as significant.

#### cDNA synthesis and qRT-PCR

DNA digestion and cDNA synthesis were performed using Turbo DNA-free DNase I kit (Ambion/Life Technologies, Darmstadt, Germany) and SuperScript III Reverse Transcriptase kit (Invitrogen/Life Technologies, Darmstadt, Germany), respectively. The qRT-PCR measurements were performed in triplicates using SYBR Green-PCR Master Mix (Applied Biosystems/Life Technologies, Darmstadt, Germany). Expression values of analyzed genes were presented in graphs as mRNA fold change. Fold change was calculated by the log_2_-normalized ΔCT to the maximum value of control treatment. The primer sequences for the reference genes and selected genes analyzed are listed in *SI Appendix*, Table S5.

#### Toluidine blue staining and RNA *in situ* hybridization

The apices of Col-0 plants grown in LD condition were harvested into formaldehyde/acetic acid/ethanol (FAA) fixative solution at 0.5, 2, 4, 24, 48, 72, 78, 96, 120, 144, 168, 192 and 216 hours after the priming (time after priming, TAP, 1^st^ stimulus) and at 0.5, 6, 24, 48, 72, 96, 120 and 144 hours after triggering (time after triggering, TAT, 2^nd^ stimulus) treatments. The following time points after priming correspond to time points after triggering: 72TAP (0.5TAT), 78TAP (6TAT), 96TAP (24TAT), 120TAP (48TAT), 144TAP (72TAT), 168TAP (96TAT), 192 TAP (120TAT), and 216TAP (144TAT) (see Fig. 1*A*). In addition, the meristems of the *hsfa2-1* mutant were harvested at 0.5 and 2h after priming, and at 0.5h after triggering treatments. After harvesting, the apices were fixed, embedded into wax using an automated tissue processor (Leica ASP200S, Leica, Wetzlar, Germany) and an embedding system (HistoCore Arcadia, Leica). Sections of 8μm thickness were prepared using a rotary microtome (Leica RM2255; Leica). Briefly, toluidine blue staining was carried out by dewaxing the slides containing longitudinal sections through apices with Histoclear and an ethanol series: 100% EtOH for 2 min, 100% EtOH for 2 min, 95% of EtOH for 1 min, 90% of EtOH for 1 min, 80% EtOH for 1 min, 60% EtOH + 0.75% of NaCl for 1 min, 30% EtOH + 0.75% of NaCl for 1 min, 0.75% NaCl for 1 min, and phosphate-buffered saline (PBS) for 1 min. After dewaxing, slides were shortly left to dry at 42°C and then incubated in 0.01% toluidine blue/sodium borate solution for 2 min, and then briefly washed with water and 80% EtOH. The sections were imaged with a Nikon Eclipse E600 microscope using NIS-Elements BR 4.51.00 software.

For RNA *in situ* hybridization, slides with sections through apices, roots, hypocotyls were washed with Histoclear solution (Biozym Scientific, Hessisch Oldendorf, Germany), and ethanol series, and Proteinase K (Roche, Mannheim, Germany). Slides were hybridized with selected probes overnight. Probes were generated from their cDNAs cloned into pGEM-T Easy Vector (Promega, Madison, Wisconsin, USA; oligo sequences are provided in Table S5) and synthesized using DIG RNA Labeling Kit (Roche). Afterwards, slides were washed out and incubated with 1% blocking reagent (Roche) in 1xTBS/0.1% Triton X-100. For immunological detection, the anti-DIG antibody (Roche) solution, diluted 1:1,250 in blocking reagent, was applied to the slides. For colorimetric detection, the NBT/BCIP stock solution (Roche), diluted 1:50 in 10% polyvinyl alcohol (PVA) in TNM-50, was applied to the slides. The slides were incubated overnight and imaged as described above. Figure panels were generated in Adobe Photoshop CS5 and Illustrator CC (Adobe Systems, San Jose, USA).

#### Iodine staining and metabolite measurements

For iodine staining, whole seedlings of Col-0 plants were harvested to 80% ethanol and boiled for 10 min, then incubated in an iodine solution (50% (v:v) Lugol’s solution) for 10 min. Excess solution was removed by washing the seedlings in water. Soluble sugars and starch content were measured in Col-0 seedlings in three biological replicates (*n=*3). Briefly, glucose, fructose, and sucrose were determined enzymatically from ethanolic extract as described (13). Starch was assayed enzymatically using pellet material (14).

#### Statistical analysis

Statistical significance between treatments was calculated using two-tailed, two-sample equal variance Student’s *t*-test: \**P*≤ 0.05; \*\**P*≤ 0.01; \*\*\**P*≤ 0.001 (Figs. 1, 3, 4, 5, 6 and *SI Appendix*, Figs. S1, S4, S8). For testing the statistical difference of RNA-seq derived gene expression levels, adjusted *P*-values were calculated by DESeq2 and edgeR with the Benjamini-Hochberg (BH) procedure for multiple testing correction (\**P*≤ 0.05) with the additional criterion of a ≥2-fold up-/down-regulation (Figs. 3 and 6 and *SI Appendix*, Figs. S4, S7, S8). Statistical significance of the enrichment of HSE motifs in 5’ regulatory regions of memory genes was calculated using the hypergeometric test compared to the regulatory regions of all TAIR10 annotated genes using the basic HSE (5’-nGAAnnTTCn-3’) and perfect HSE definition (5’-GAAnnTTCnnGAA-3’).

## Supplementary Figures

**Fig. S1.**
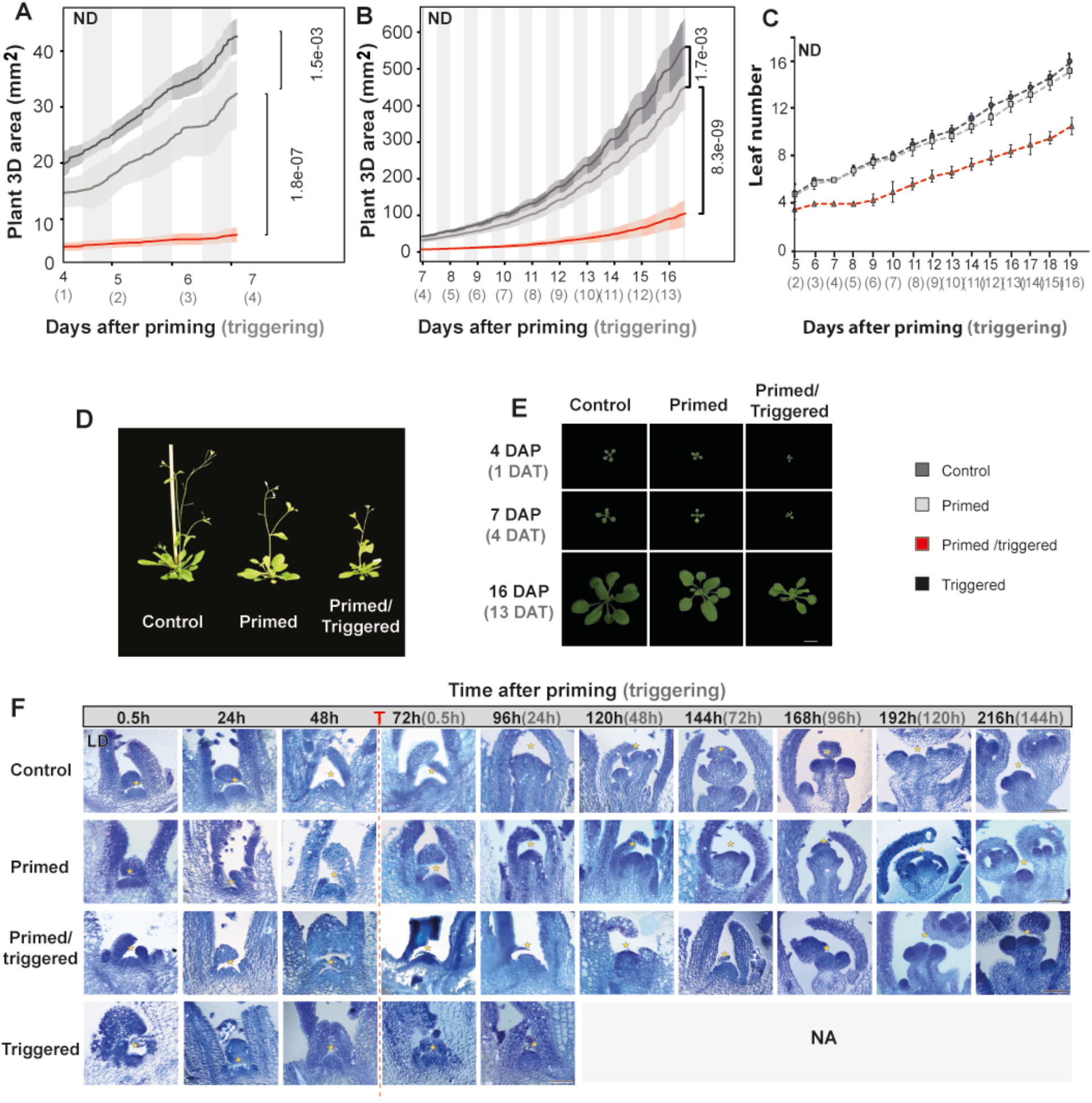
Morphological analyses of wild-type (Col-0) plants grown in long day (LD) and neutral (ND) conditions during and after thermopriming. *(A, B)* Increase of total 3D area over time of control (C), primed (P) and primed and triggered (PT) plants grown in ND photoperiod, analysed using the Phenotyping^4D^ platform *(A, B)*. Note, seedlings that only obtained the triggering (T) stimulus died. *(C)* Leaf initiation rate analyzed in ND conditions determined by counting the appearance of 2mm-sized leaves throughout vegetative development. *(D)* Flowering time phenotype of Col-0 plants after thermopriming. *(E)* Images of Col-0 plants after priming (DAP) and triggering (DAT) treatments. *(F)* Toluidine blue-stained longitudinal sections through apices of C, P, PT and T plants after thermopriming in LD. Note, morphological analysis of the meristem of T plants was performed until 96h after priming (24h after triggering). Due to lethality of the plants further time points were not analyzed (NA). Time is given in hours (h) after priming (black color) and triggering (grey color) treatment. The vertical dashed line represents the time point of triggering (T) treatment. Error bars indicate s.d. Asterisks indicate meristem summit. Scale bars, 1cm *(E)* and 100µm *(F)*.

**Fig. S2.**
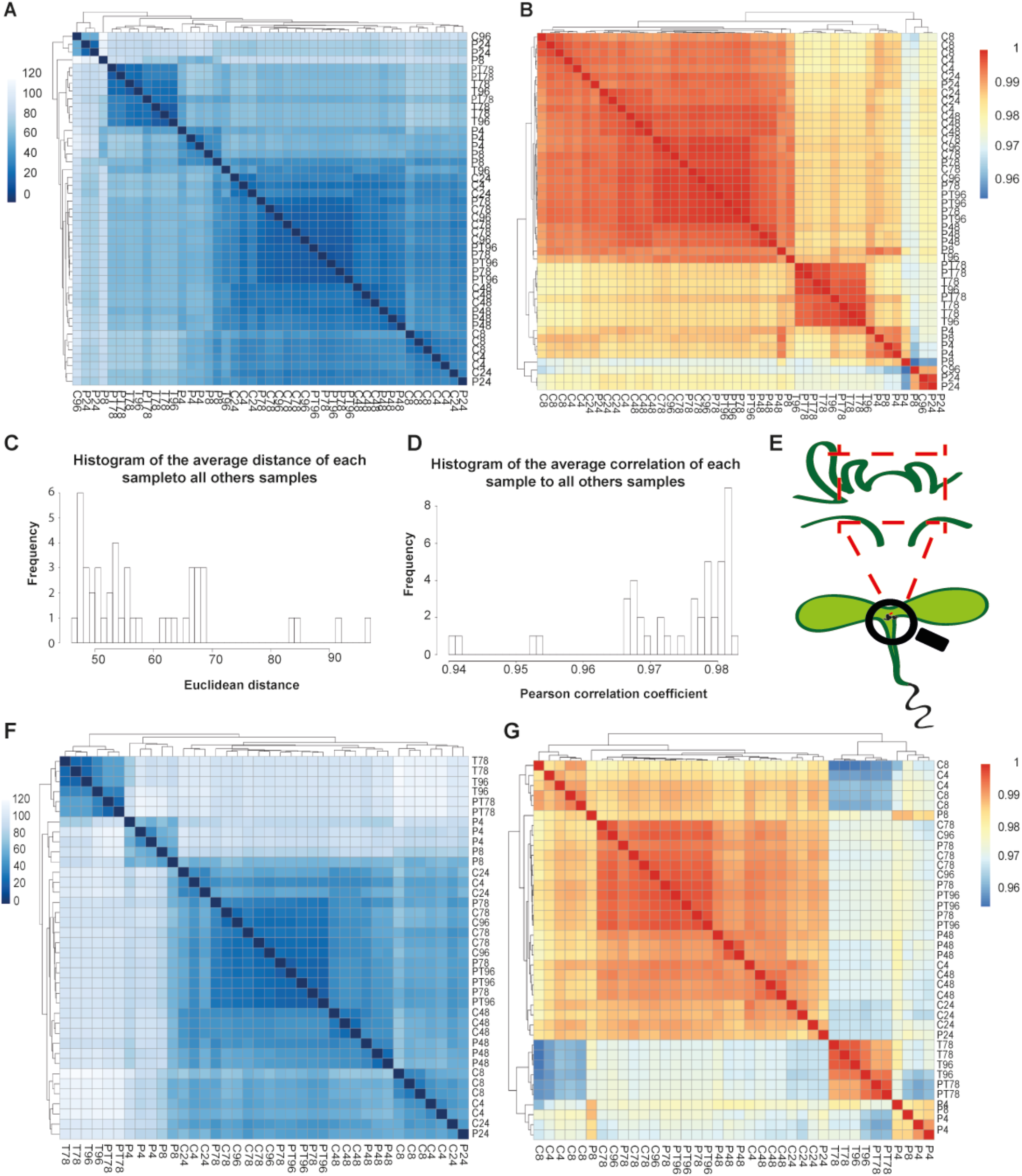
Clustering of gene expression patterns of all 45 samples after variance stabilizing transformation (VST) of the DESeq2 package. *(A)* Heatmap of the distance matrix of all 45 samples using pairwise Euclidean distance. *(B)* Heatmap of the correlation matrix of all 45 using pairwise Pearson correlation. *(C)* Histogram of the average distance of each sample to other samples. *(D)* Histogram of the average correlation of each sample to all other samples. Note, both clustering approaches revealed four outlier samples (sample degradation or/and low number of reads) with an average Euclidean distance of ≥80 and a Pearson correlation value of <0.96; those samples were removed for further analysis (1x P8, 2x P24, 1x C96). Furthermore, to increase the power of the differential gene expression (DE) analysis, for each triplicate we filtered samples whose average Euclidean distance to the remaining two triplicates is more than 50% higher than the distance of the other two replicates to each other (1x PT78, 1x T78, 1x T96). Thus, the filtered dataset contains 38 samples from 15 different experimental conditions with triplicates or duplicates with the exception of P24, which remains a single sample that makes DE analysis at 24h after priming not feasible, however, allows representation without standard deviation in time-course plots (see e.g. Fig. 3, and *SI Appendix*, Fig, S4 and S8). *(E)* Schematic representation of the material harvested and used for RNA-seq analysis. *(F)* Heatmap of the distance matrix of 38 samples using pairwise Euclidean distance. *(B)* Heatmap of the correlation matrix of 38 samples using pairwise Pearson correlation.

**Fig. S3.**
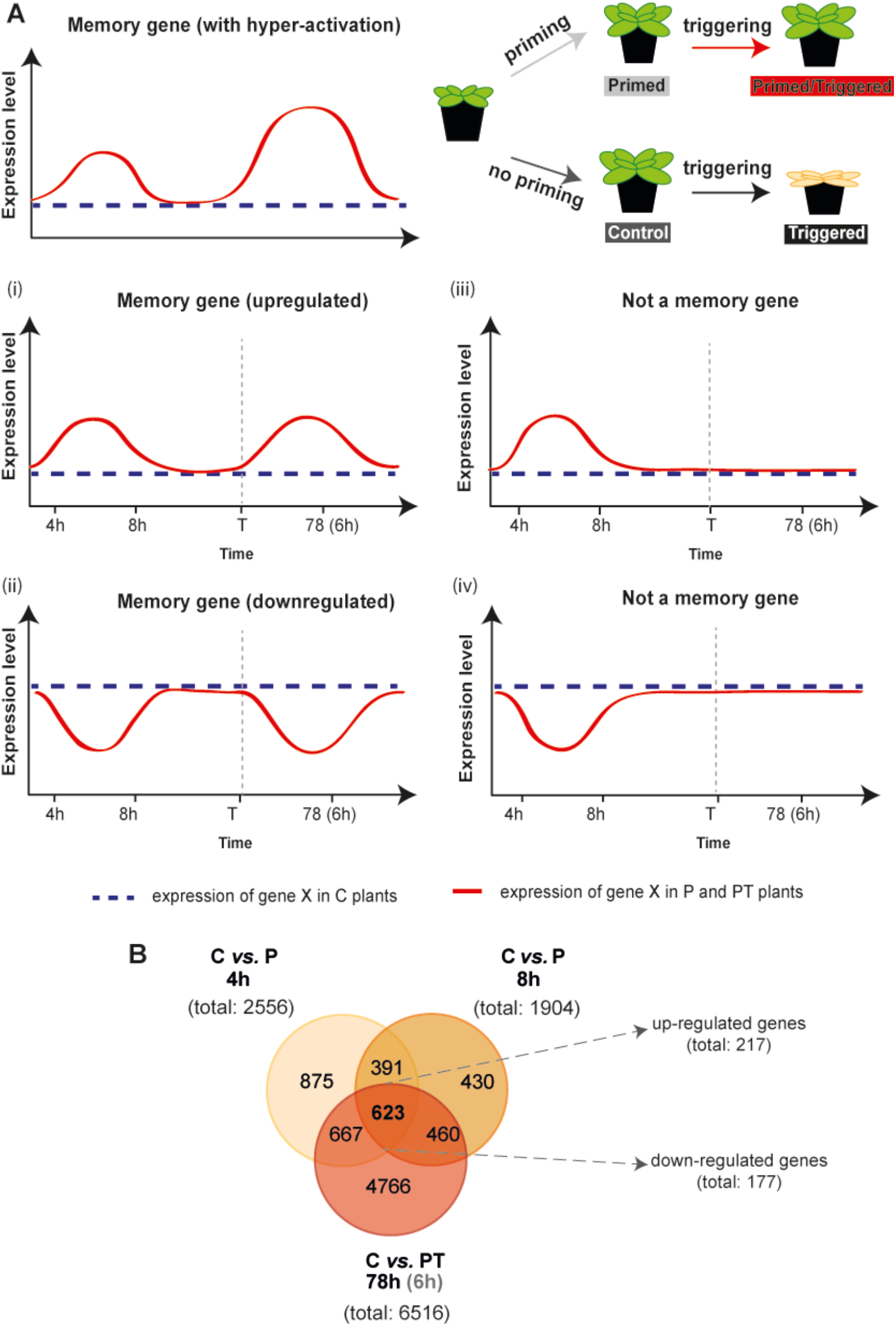
Identification of thermomemory genes at the shoot apex. *(A)* Transcriptional memory genes were identified by RNA-seq as: (i) genes whose expression level was significantly upregulated at 4 and 8h after priming and at 6h after triggering (72h after priming) compared to control (C) condition, and (ii) genes whose expression level was significantly downregulated at 4 and 8h after priming and at 6h after triggering (72h after priming) compared to C condition. Note, that genes whose expression was induced (iii) or downregulated (iv) only by the priming stimulus were not considered as memory genes. *(B)* Venn diagram of DEGs at 4h and 8h after priming and 6h after triggering (78h after priming) of primed (P) and primed and triggered (PT) plants compared to the control (C). The overlap represents significantly changed HS memory genes at the shoot apex of Col-0 plants during thermopriming.

**Fig. S4.**
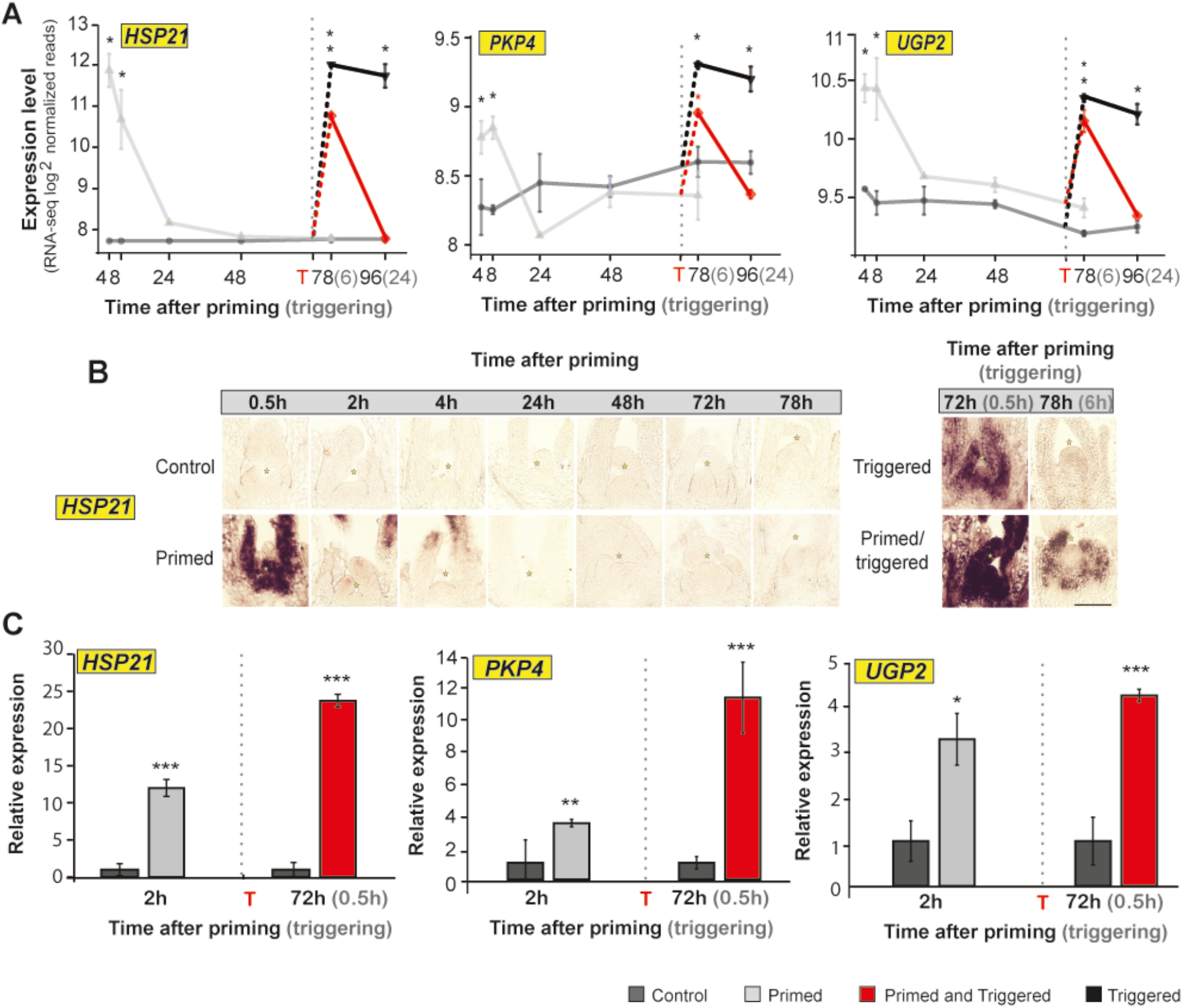
Expression of HS memory-induced genes at the shoot apical meristem (SAM) of Col-0 wild-type plants. *(A)* Relative expression level of *HEAT SHOCK PROTEIN 21* (*HSP21*), *PYRUVATE KINASE 4* (*PKP4*) and *UDP-GLUCOSE PYROPHOSPHORYLASE 2 (UGP2)* genes at the shoot apex of Col-0 plants obtained by RNA-seq (*n*=3). *(B)* RNA-seq results by RNA *in situ* hybridization using *HSP21* as probe on longitudinal section through meristems of Col-0 plants. Scale bars, 100µm. *(C)* Expression level of *HSP21, PFP4* and *UGP2*, at the SAM of Col-0 plants during thermopriming obtained by qRT-PCR. Time is given in hours (h) after priming (black color) and triggering (grey color) treatments. The vertical dashed line represents the time point of triggering (T) treatment. Error bars indicate s.d. (*n*=3). Asterisks indicate meristem summit *(B)* or statistically significant difference ((*A)* \**P*≤ 0.05 adjusted with Benjamini-Hochberg procedure for multiple testing correction; *(C)* Student’s t-test: \**P*≤ 0.05; \*\**P*≤ 0.01; \*\*\**P*≤ 0.001) from the control conditions.

**Fig. S5.**
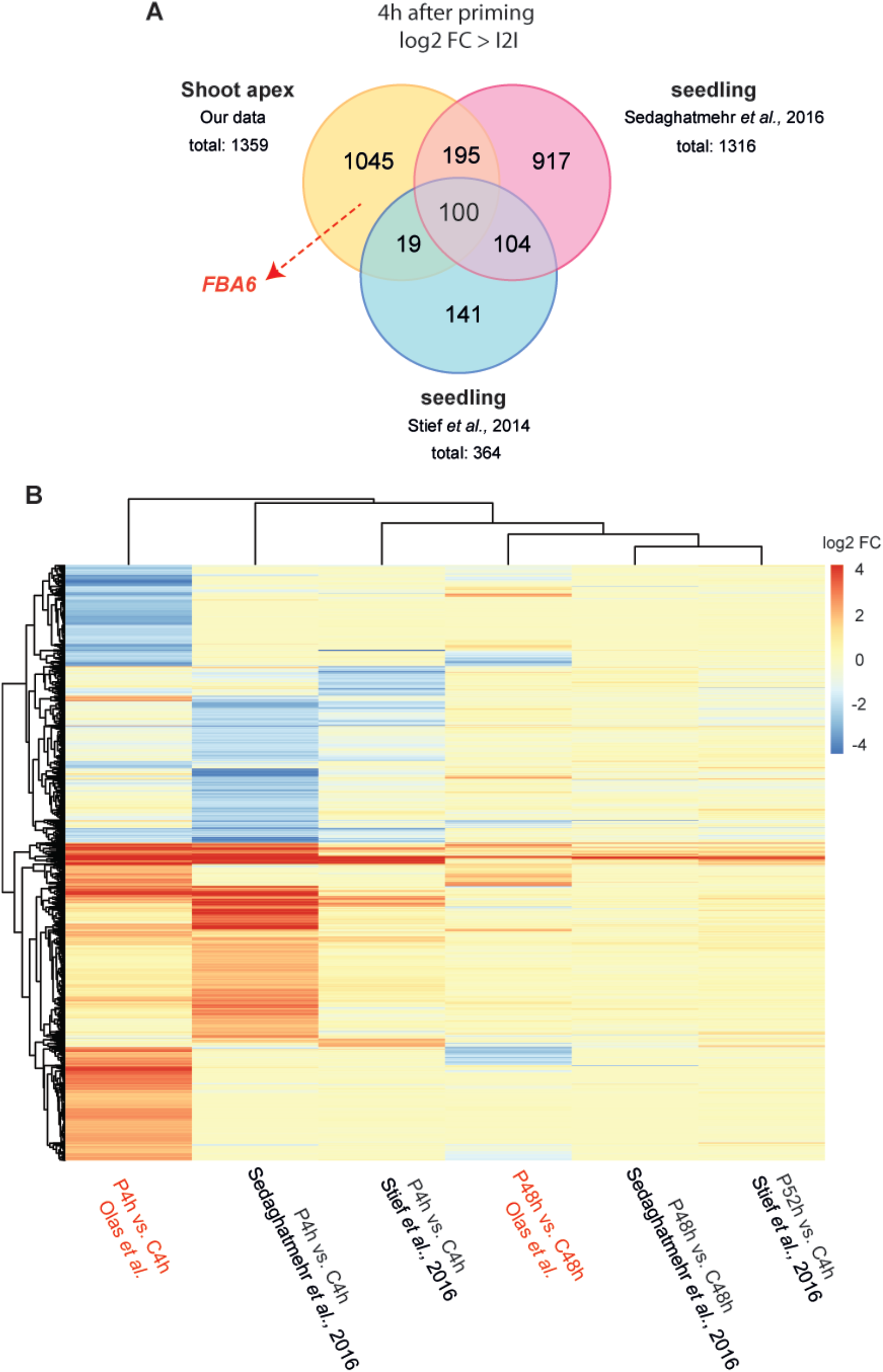
Tissue-dependent variation in the regulation of thermomemory. *(A)* Venn diagram representation of fold change expression of genes identified at the shoot apex and in whole seedlings of Arabidopsis (Stief *et al*., 2014; Sedaghatmehr *et al*., 2016) at 4h after priming with log_2_FC>|2|. Note, that priming treatment was performed in the same way in all studies. *(B)* Heat map visualizing the responses of genes changed at 4h and 48/52h after priming (priming (P) *versus* control (C); log_2_FC>|2|; 2,521 genes) between shoot apex and whole seedlings of Arabidopsis (Stief *et al*., 2014; Sedaghatmehr *et al*., 2016) with log_2_FC>|2|. Note, data reported by Stief *et al*. (2014) and Sedaghatmehr *et al*. (2016) were obtained by microarray analyses. The published data was downloaded from NCBI GEO and processed to obtain log_2_FC values compared to control allowing comparison to the RNA-seq data.

**Fig. S6.**
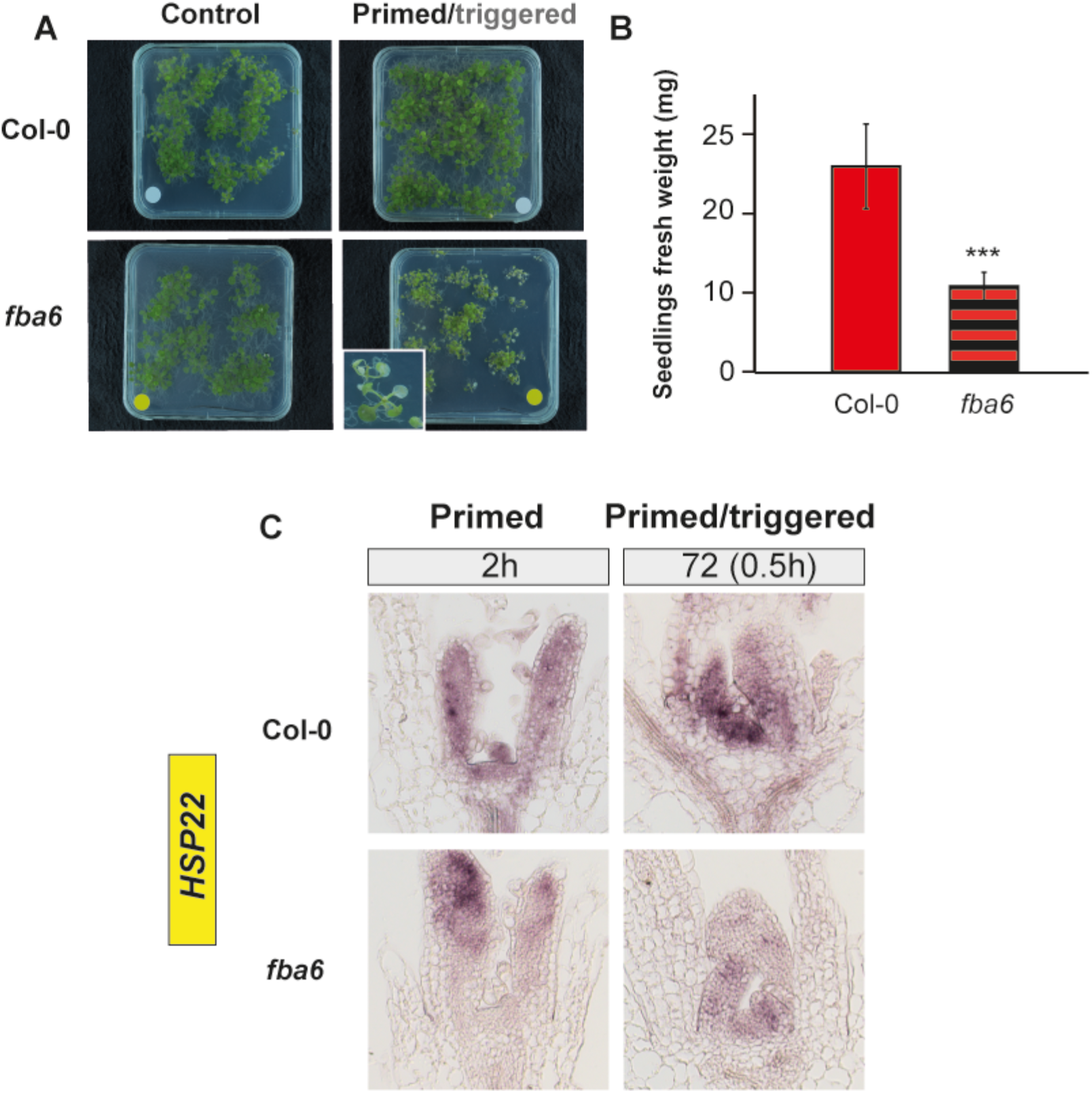
Growth recovery of *fba6* and *fba8* mutants and Col-0 wild-type plants. *(A)* Phenotype of Col-0 and *fba6* seedlings 5 days after triggering. *(B)* Fresh weight of results shown in (A) for PT Col-0 and *fba6* plants. Error bars indicate s.d. (*n*>12). Asterisks indicate statistically significant difference (Student ‘s *t*-test: ***P≤ 0.001) from Col-0. *(C)* RNA *in situ* hybridization on longitudinal sections through apices of primed and primed/triggered Col-0 wild-type and *fba6* mutant plants using a specific probe against *HEAT SHOCK PROTEIN 22* (*HSP22*).

**Fig. S7.**
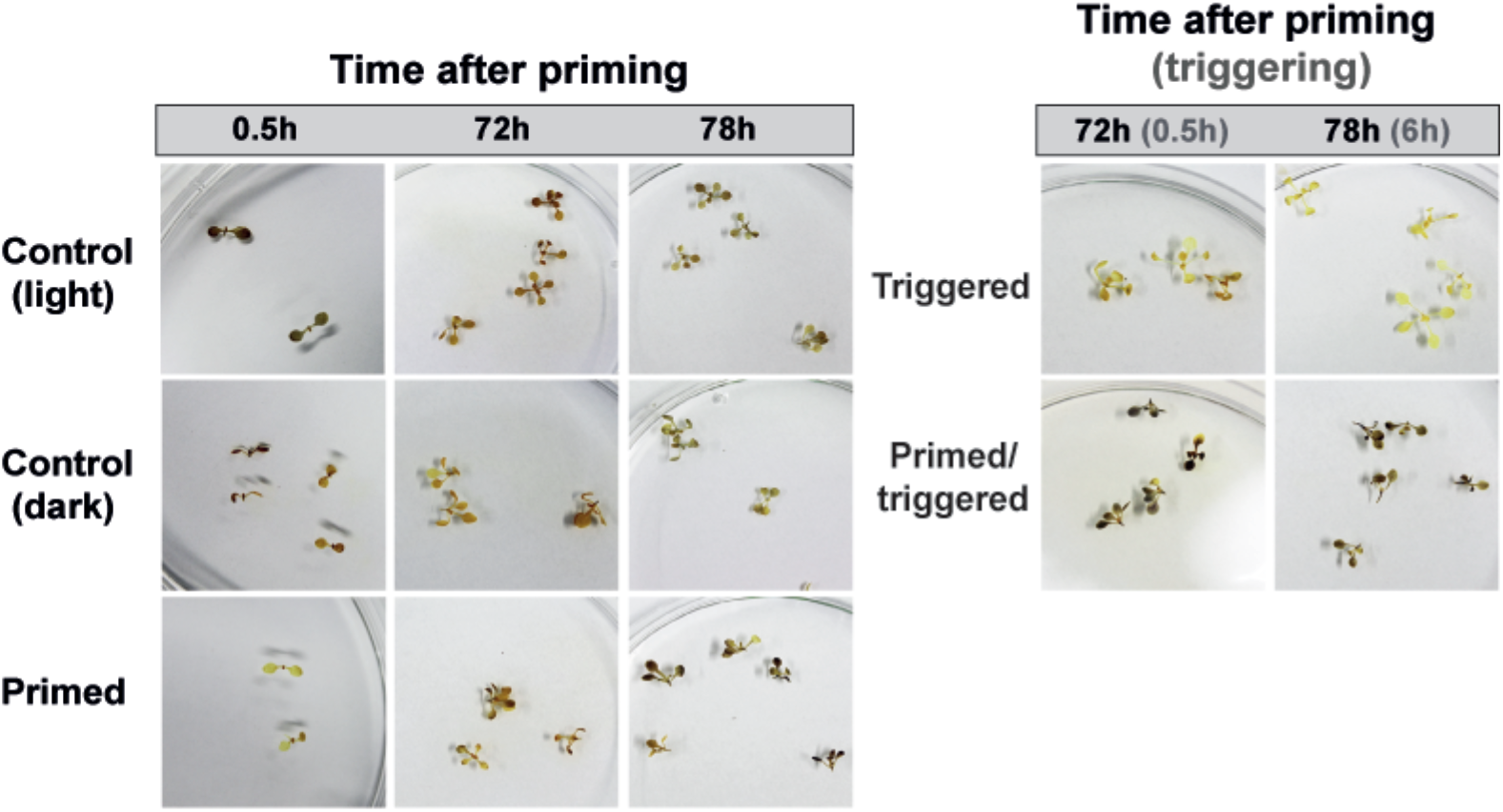
Sugar availability. Iodine staining of Col-0 wild-type plants during thermopriming. Time is given in hours (h) after priming (black color) and triggering (grey color) treatments.

**Fig. S8.**
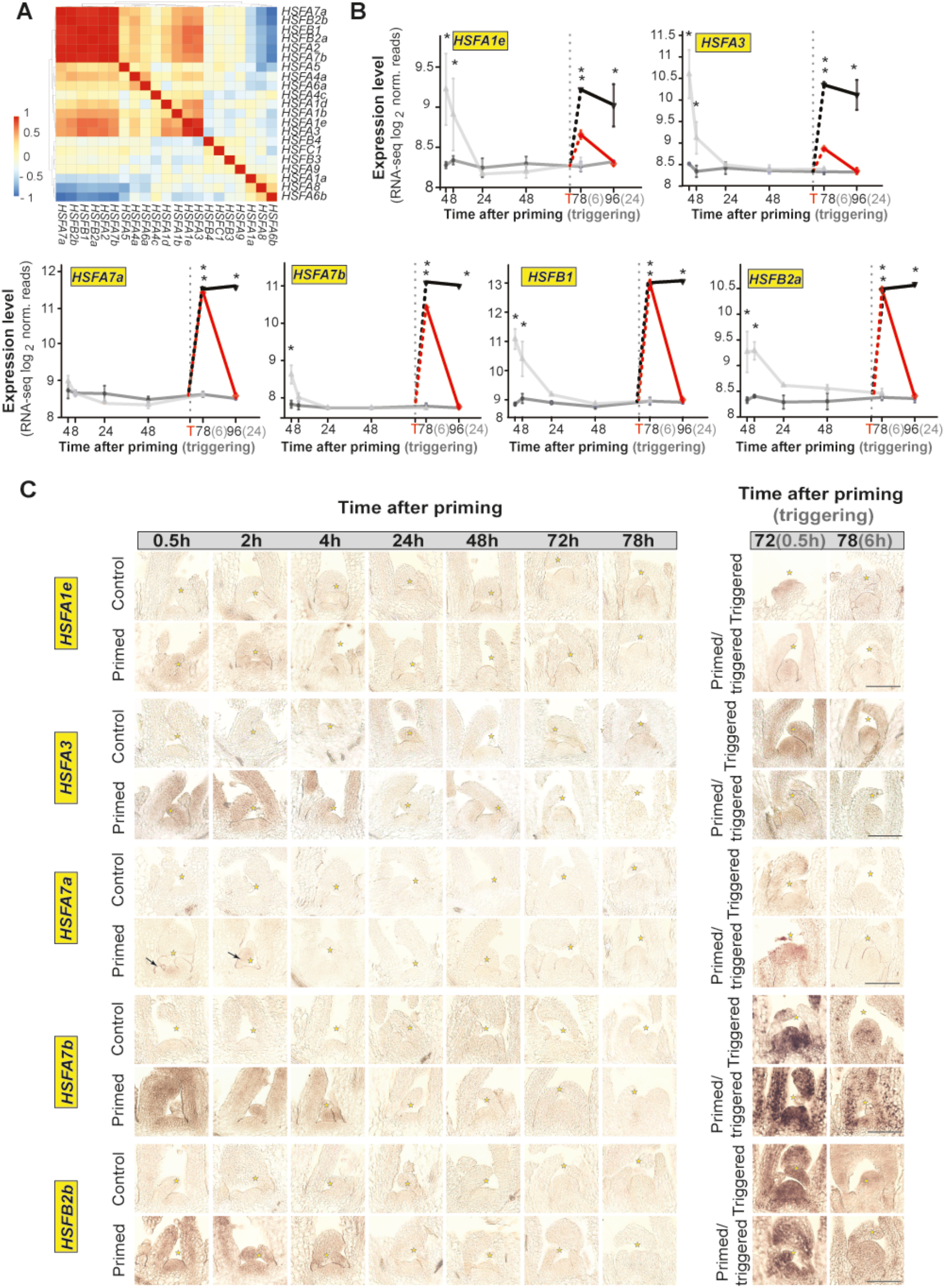
*HSFs* expressed in the shoot apex. *(A)* Clustering of *HEAT SHOCK TRANSCRIPTION FACTORs* (*HSF*s) using correlation as distance measure. *(B)* Expression level of induced *HSFs* in the shoot apex analyzed by RNA-seq. Error bars indicate s.d. (*n*=3). Asterisks indicate statistically significant difference (\**P*<0.05 adjusted with Benjamini-Hochberg procedure for multiple testing correction) from the control conditions. The vertical dashed line represents the time point of triggering (T) treatment. *(C)* RNA *in situ* hybridization on longitudinal sections through apices of control, primed, primed and triggered and triggered seedlings grown in LD condition using specific probes against *HSFs*. Asterisks indicate meristem summit. Scale bars, 100µm. Time is given in hours (h) after priming (black color) and triggering (grey color) treatment.

**Fig. S9.**
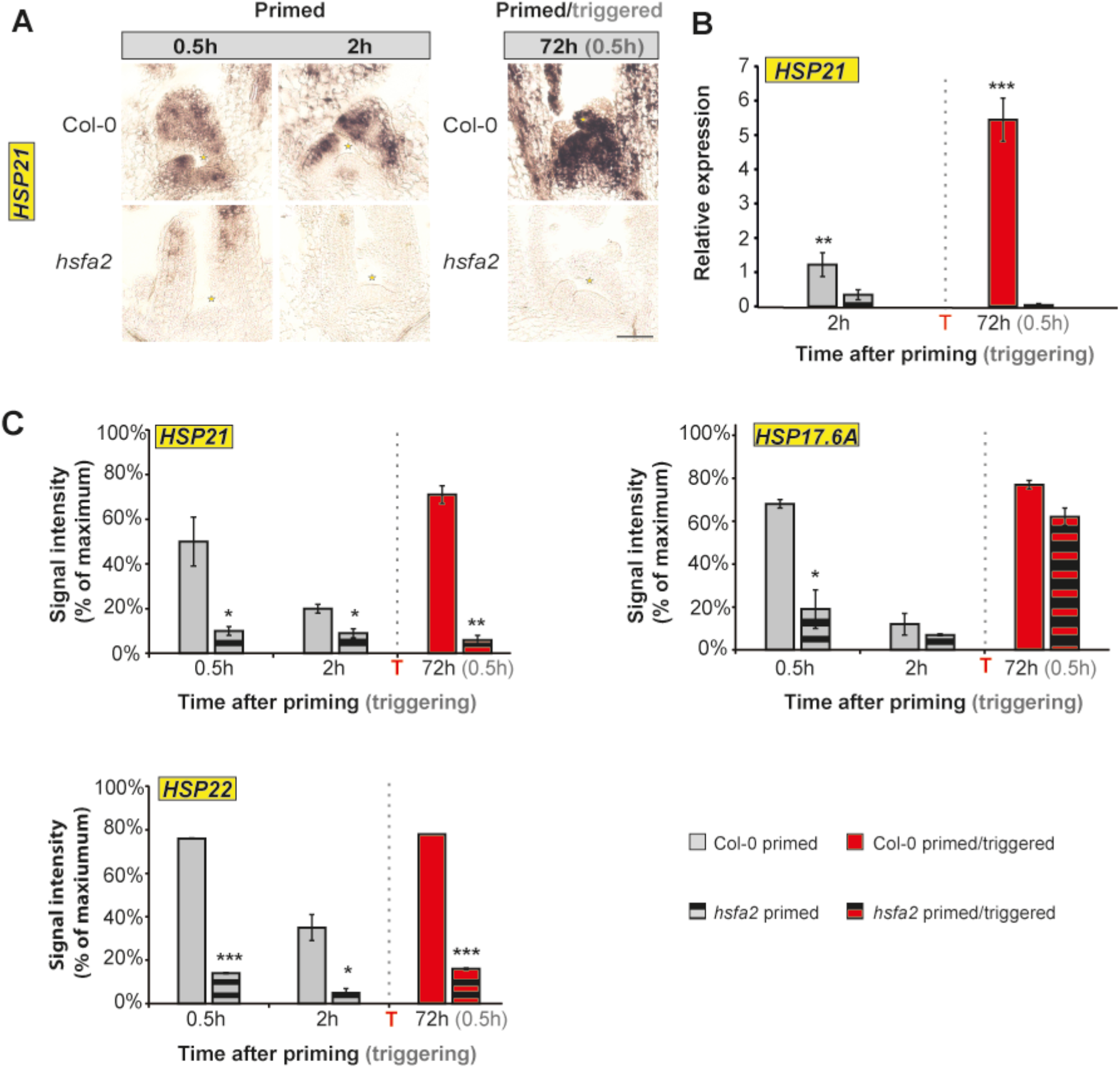
Expression of *HSPs* at the SAM of Col-0 and *hsfa2* mutant plants. *(A)* RNA *in situ* hybridization on longitudinal sections through apices of Col-0 wild-type and *hsfa2* mutant plants using a specific probe against *HEAT SHOCK PROTEIN 21* (*HSP21*). Scale bars, 100µm. *(B)* Expression level of *HSP21* at the SAM of Col-0 wild-type and *hsfa2* mutants analyzed by qRT-PCR. *(C)* Signal intensity (% of maximum) of *HSP21, HSP17*.*6A* and *HSP22* measured at the SAM of Col-0 and *hsfa2* mutant plants. Error bars indicate s.d. (*n*=3). Asterisks indicate meristem summit *(A)* or statistically significant difference (Student’s *t*-test: \**P*≤0.05; \*\**P*≤ 0.01; \*\*\**P*≤ 0.001; *(B, C)*) from control conditions. Time is given in hours (h) after priming (black color) and triggering (grey color) treatment. The vertical dashed line represents the time point of the triggering (T) treatment.

**Table S1.** Flowering time data described in this study.

| Treatment (Col-0; long days) | DTB | TLN | LIR | n |
| --- | --- | --- | --- | --- |
| Control | **19.3 ± 0.5** | 10.7 ± 1.0 | 0.54 ± 0.06 | 13 |
| Primed | **21.4 ± 1.6<sup>+</sup>** | 9.1 ± 0.6<sup>+</sup> | 0.43 ± 0.05<sup>+</sup> | 16 |
| Primed/triggered | **24.1 ± 2.2<sup>+</sup>** | 11.7 ± 1.3<sup>+</sup> | 0.49 ± 0.1<sup>+</sup> | 15 |
| Triggered | n.a. | n.a. | n.a. | n.a. |

Data represent averages of at least 20 genetically identical replicate plants. Abbreviations: DTB, days to bolting, in bold as referred to in the main text; TLN, total leaf number; LIR, leaf initiation rate; n, number of individuals; (+/-), presence or absence of significance based on Student's *t*-test calculated between control and treated plants, respectively (+: *P*-value ≤ 0.05, -: *P*-value ≥ 0.05); n.a., not analysed.

**Table S2.** Number of significantly changed genes with and without log_2_ FC > |1|. Number of upregulated and downregulated genes is indicated in brackets.

|  | 4h | 8h | 48h | 78h | 96h |
| --- | --- | --- | --- | --- | --- |
| **DEGs:significant** |  |  |  |  |  |
| **C versus P** | 2556 (1259;1297) | 1904 (968;936) | 1241 (549;692) | 51 (8;43) | - |
| **C versus PT** |  |  |  | 6516 (3658;2858) | 120 (40;80) |
| **C versus T** |  |  |  | 6108 (3574;2534) | 3298 (2046;1252) |
| **PT versus T** |  |  |  | 2496 (1334;1162) | 6439 (3706;2733) |
| **DEGs: significant and $\log_2 \text{FC} > \vert 1\vert $** |  |  |  |  |  |
| **C versus P** | 1175 (664;511) | 780 (404, 376) | 203 (106;97) | 0 (0;0) | - |
| **C versus PT** |  |  |  | 2105 (1050;1055) | 1 (0;1) |
| **C versus T** |  |  |  | 2054 (1202;852) | 1634 (1053;581) |
| **PT versus T** |  |  |  | 631 (441;190) | 2219 (1300;916) |

Abbreviations: C, control; P, primed; PT, primed and triggered; T, triggered.

**Table S3.**
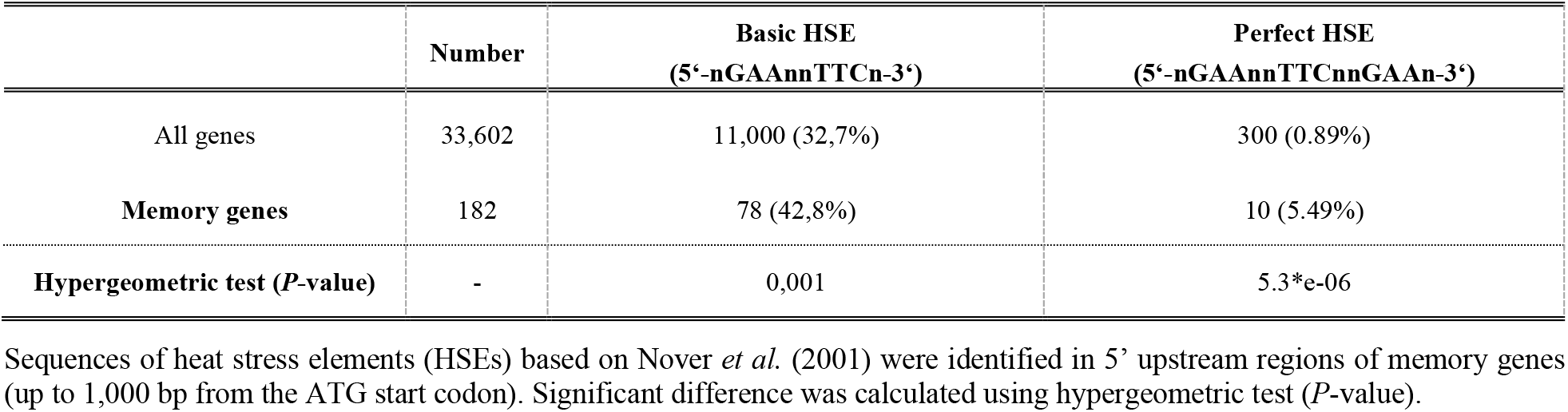
Analysis of 5’ upstream regulatory regions of identified memory genes.

**Table S4.** List of memory genes containing HSE in 5’ upstream regulatory regions.

| ATG Identifier | Symbol | Basic HSE | Perfect HSE |
| --- | --- | --- | --- |
| AT1G03070 | LFG4 | + |  |
| AT1G04130 | TPR2 | + |  |
| AT1G21550 | AT1G21550 | + |  |
| AT1G26800 | MPSR1 | + |  |
| AT1G30070 | AT1G30070 | + |  |
| AT1G48710 | Transposable element gene | + |  |
| AT1G52560 | AT1G52560 | + |  |
| AT1G52870 | AT1G52870 | + |  |
| AT1G53540 | AT1G53540 | + | + |
| AT1G54050 | AT1G54050 | + |  |
| AT1G66510 | AT1G66510 | + |  |
| AT1G67360 | SRP1 | + |  |
| AT1G71000 | AT1G71000 | + | + |
| AT1G72610 | GER1 | + |  |
| AT1G73480 | MAGL4 | + |  |
| AT1G74310 | HSP101 | + |  |
| AT1G79920 | HSP70-15 | + |  |
| AT2G13550 | AT2G13550 | + |  |
| AT2G20560 | AT2G20560 | + |  |
| AT2G21820 | AT2G21820 | + |  |
| AT2G26150 | HSFA2 | + | + |
| AT2G29500 | AT2G29500 | + |  |
| AT2G32120 | HSP70T-2 | + |  |
| AT2G32860 | BGLU33 | + |  |
| AT2G36460 | FBA6 | + |  |
| AT2G46240 | BAG6 | + |  |
| AT2G47180 | GOLS1 | + |  |
| AT3G02990 | HSFA1E | + |  |
| AT3G03270 | HRU1 | + |  |
| AT3G04720 | PR4 | + |  |
| AT3G07150 | AT3G07150 | + |  |
| AT3G08690 | UBC11 | + |  |
| AT3G08770 | LTP6 | + |  |
| AT3G08970 | TMS1 | + |  |
| AT3G09350 | FES1A | + |  |
| AT3G09640 | APX2 | + |  |
| AT3G10020 | AT3G10020 | + |  |
| AT3G12145 | FLOR1 | + |  |
| AT3G12580 | HSP70 | + | + |
| AT3G15770 | AT3G15770 | + |  |
| AT3G16530 | AT3G16530 | + |  |
| AT3G22840 | ELIP1 | + | + |
| AT3G24100 | AT3G24100 | + |  |
| AT3G24500 | MBF1C | + |  |
| AT3G46230 | HSP17.4 | + | + |
| AT3G63310 | LFG2 | + |  |
| AT4G04020 | FIB1A | + |  |
| AT4G10040 | CYTC-2 | + |  |
| AT4G17250 | AT4G17250 | + |  |
| AT4G21323 | AT4G21323 | + |  |
| AT4G23493 | AT4G23493 | + |  |
| AT4G23680 | AT4G23680 | + |  |
| AT4G25200 | ATHSP23.6-MITO | + |  |
| AT4G25810 | XTR6 | + |  |
| AT4G27670 | HSP21 | + |  |
| AT5G03340 | ATCDC48C | + |  |
| AT5G05410 | DREB2A | + |  |
| AT5G12020 | HSP17.6II | + |  |
| AT5G13200 | GER5 | + |  |
| AT5G17310 | UGP2 | + |  |
| AT5G25450 | AT5G25450 | + |  |
| AT5G50240 | PIMT2 | + |  |
| AT5G51440 | AT5G51440 | + |  |
| AT5G51740 | OMA1 | + |  |
| AT5G52640 | HSP90.1 | + | + |
| AT5G58770 | CPT7 | + |  |
| AT5G59720 | HSP18.2 | + | + |
| AT5G59820 | ZAT12 | + | + |
| AT5G62020 | HSFB2A | + | + |
| AT5G64170 | LNK1 | + |  |
| AT5G64510 | TIN1 | + |  |
| ATMG00160 | COX2 | + |  |
| ATMG00516 | NAD1C | + |  |
| ATMG00670 | ORF275 | + |  |
| ATMG01120 | NAD1B | + |  |
| ATMG01130 | ORF106F | + |  |
| ATMG01360 | COX1 | + |  |

**Table S5.**
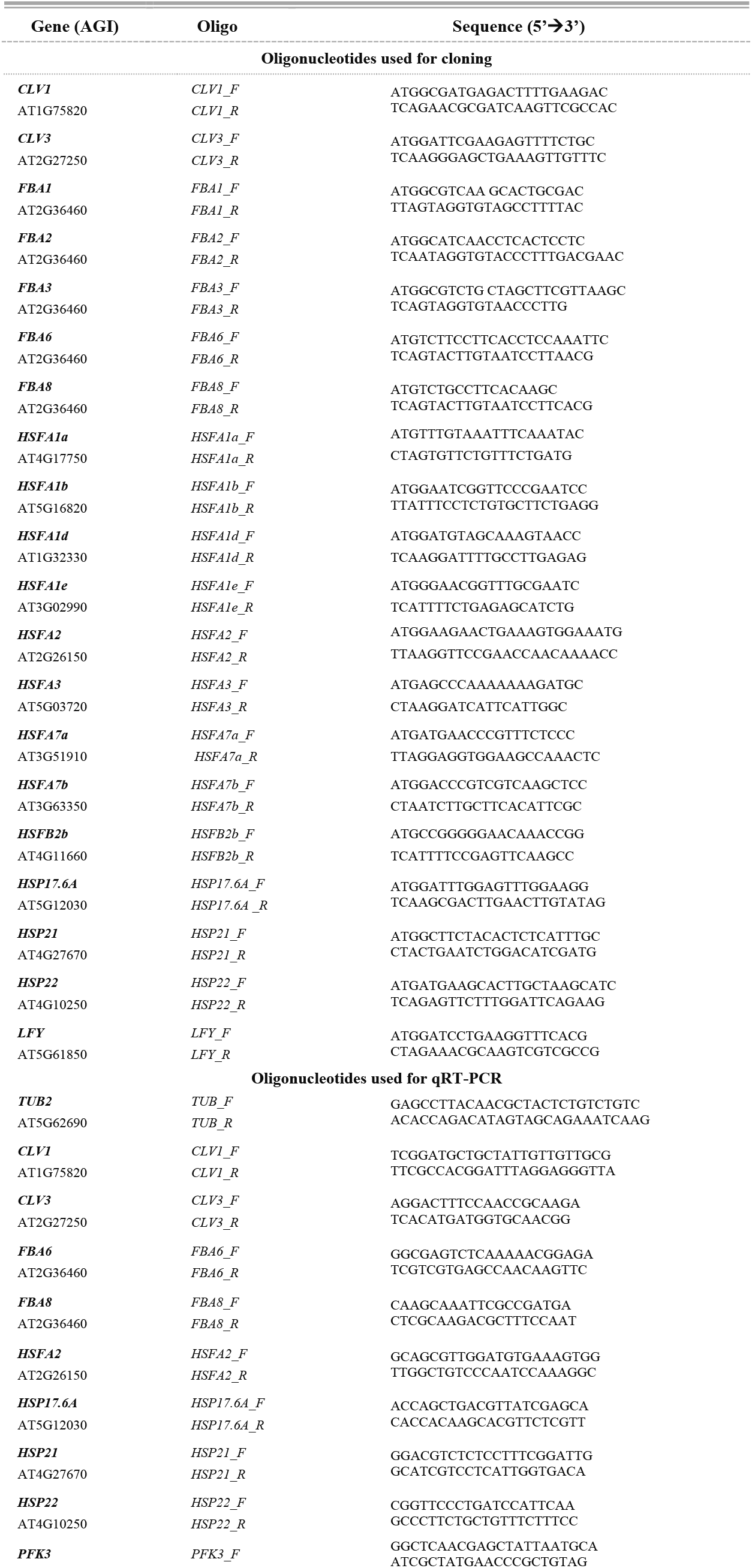
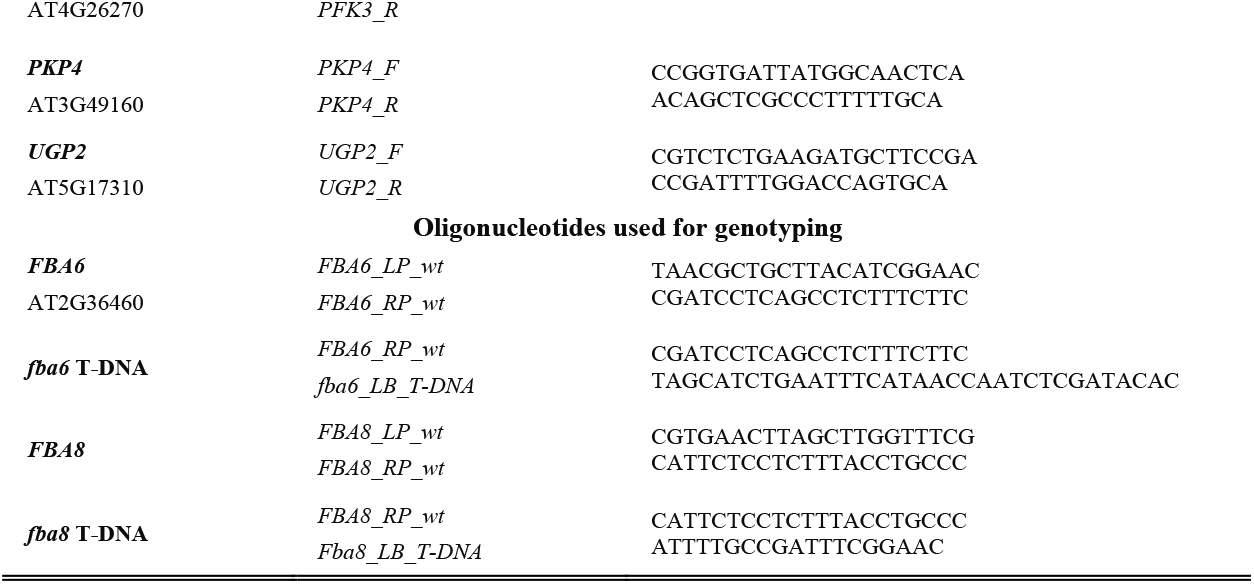
Oligonucleotides used in this study.

